# Human growth plates house resting zone sub-populations with features of quiescent stem cells

**DOI:** 10.1101/2025.03.12.642613

**Authors:** Mahtab Avijgan, Ana R.L. Perez, Leire A. Galicia, Laura Sudupe, Jose G. Marchan-Alvarez, Amal Nazaraliyev, Žaneta Andrusivová, Ludvig Larsson, Yunhan Zhao, Farasat Zaman, Hong Qian, Klas Blomgren, Felipe Prosper, Jesper N. Tegner, Joakim Lundeberg, Lars Sävendahl, Reza Mirzazadeh, David Gomez-Cabrero, Phillip T. Newton

**Author notes:** These authors contributed equally to this work (Mahtab Avijgan, Ana R.L. Perez, Leire A. Galicia). These authors contributed equally to this work (Reza Mirzazadeh, David Gomez-Cabrero, Phillip T. Newton).

## Abstract

Incomplete mapping of gene expression within human (epiphyseal) growth plates contributes to the challenges of diagnosing and treating patients with skeletal growth disorders. To address this issue, we applied spatially resolved transcriptomics to rare growth plate biopsies obtained from healthy adolescents. In addition to identifying novel markers of each zone of the human growth plate, spatial profiling revealed that the expression of genes associated with poorly understood growth disorders, including NKX3-2, SGMS2 and WNK4, is restricted to specific human growth plate zones. By elaborating on the low transcriptional activity of resting zone chondrocytes, we found that a subset of these cells exists in a functionally quiescent state *in vivo*, as determined by their predominantly nuclear mRNA, abundant heterochromatin, and ability to exit the G0 phase under specific conditions - features shared with skeletal stem cells in mouse growth plates. Additionally, we identified distinct and overlapping sub-populations of human resting zone chondrocytes; an exploration of their hierarchy determined that CHRDL2 and/or SFRP5-positive sub-populations are among the least quiescent resting zone cells. In summary, we generated the most comprehensive gene expression characterization of the human growth plate, which revealed novel zone-specific markers, new primary growth disorders, candidate pharmacological targets, and led us to uncover sub-populations of resting zone chondrocytes with features of quiescent stem cells. These results contribute to a better understanding of the cellular and molecular mechanisms governing human height and can facilitate improved diagnosis and treatment strategies of patients with skeletal growth disorders.

## INTRODUCTION

During childhood and adolescence, humans grow taller because tubular bones, such as femora and tibiae, increase in length. These bones are elongated by narrow cartilage discs called (epiphyseal) growth plates, which are positioned between the bony primary and secondary ossification centers (POC and SOC, respectively) (*1*). The growth plate forms once a SOC spatially separates the articular from the growth cartilage, which occurs around birth in humans (*2*). Histologically, each growth plate can be divided into three distinct zones: directly adjacent to the SOC, slowly dividing chondro-progenitors comprise the resting zone (RZ); some of these cells give rise to columns of flattened chondrocytes that undergo several cycles of cell division in the proliferating zone (PZ), and subsequently differentiate and increase in size in the hypertrophic zone (HZ), producing a mineralized matrix that is used as a scaffold for new bone tissue to be built on (*3*). Since this process of bone elongation continues until the end of puberty, its regulation is crucial for humans to reach their expected height and body proportions.

Skeletal growth disorders are categorized as primary (directly affecting growth plates), secondary (indirectly affecting growth plates, via, for example, hormones) or idiopathic (*4*). However, distinguishing primary growth disorders from secondary, and elucidating idiopathic ones remains challenging because, while candidate genes can be identified using whole exome/genome sequencing, our understanding of which genes are expressed in human growth plates is severely limited (*5*). Many growth plate markers are known only from animal experiments, and their relevance to humans remains uncertain. This knowledge gap is also reflected by the unrefined nature of therapeutic approaches: the most common treatment for patients with idiopathic short stature with growth hormone (GH) sufficiency is recombinant GH, particularly in the United States of America (*5*). For idiopathic tall stature adolescents, only invasive surgery to remove the growth plate has been proven to be a safe and effective treatment (*6*). Hence, incomplete mapping of gene expression in human growth plates contributes to a poor understanding of the mechanisms underpinning human skeletal growth and a lack of pharmacological targets to treat growth disorders with precision.

Tubular bone growth is considered to be highly evolutionarily conserved within all extant mammals (*7*). Experiments in mice suggest that once the SOC forms in the epiphysis, growth is maintained by skeletal stem cells located in the RZ of growth plates (*8*). Although growth mechanisms are similar between mice and humans, key differences between these species include gross length, bipedal versus quadrupedal posture, and the relative sizes of growth plate zones (*9*). Furthermore, humans experience finite growth, with growth plates disappearing entirely at the end of puberty - a process that does not typically occur in rodents (*10*). Understanding the cell populations within human growth plates could lead to a better understanding of human growth maintenance and more effective treatment strategies.

To better describe the human growth plate, we applied spatially resolved transcriptomics (SRT) with 10x Genomics’ Visium platform to rare samples of human growth plate sections obtained from healthy adolescents. This approach allowed us to obtain gene expression profiles while maintaining cellular positioning and growth plate architecture. We identified a panel of 138 marker genes specific to individual growth plate zones and SOC, allowing us to verify growth plate markers reported in other species, identify novel markers of each growth plate zone and uncover previously unknown primary growth disorders. By elaborating upon their low transcriptional activity, we found that RZ chondrocytes reside in a functionally quiescent state *in vivo* and have similar properties to skeletal stem cells in murine growth plates. Finally, by exploring the expression of RZ markers in association with quiescence, we reveal a potential hierarchy among sub-populations of RZ cells.

## RESULTS

### Resting zone chondrocytes contain relatively low amounts of RNA

We obtained biopsies containing human growth plates from healthy adolescents aged 12 to 14 years who were undergoing epiphysiodesis surgery for idiopathic tall stature (Fig. 1A, Table 1). Sections of the biopsies spanning the growth plate and SOC were prepared for analysis by Visium. We previously established a modified version of the 10x Genomicś Visium protocol, which we called RNA-Rescue Spatial Transcriptomics (RRST), that enabled the examination of spatial transcriptional profiles within non-decalcified cartilage (*11*). First, we applied the RRST protocol to human growth plate biopsies (Fig. 1B), using histology to manually define four epiphyseal areas: SOC, RZ, PZ and HZ. We observed that spots corresponding to cartilage had relatively low unique molecular identifier (UMI) counts (total number of transcripts captured in each spot), compared to areas of the bone containing bone marrow (Fig. 1B, C, fig. S1A), which were consistent with those expected in other tissues (*11*).

**Figure 1.**
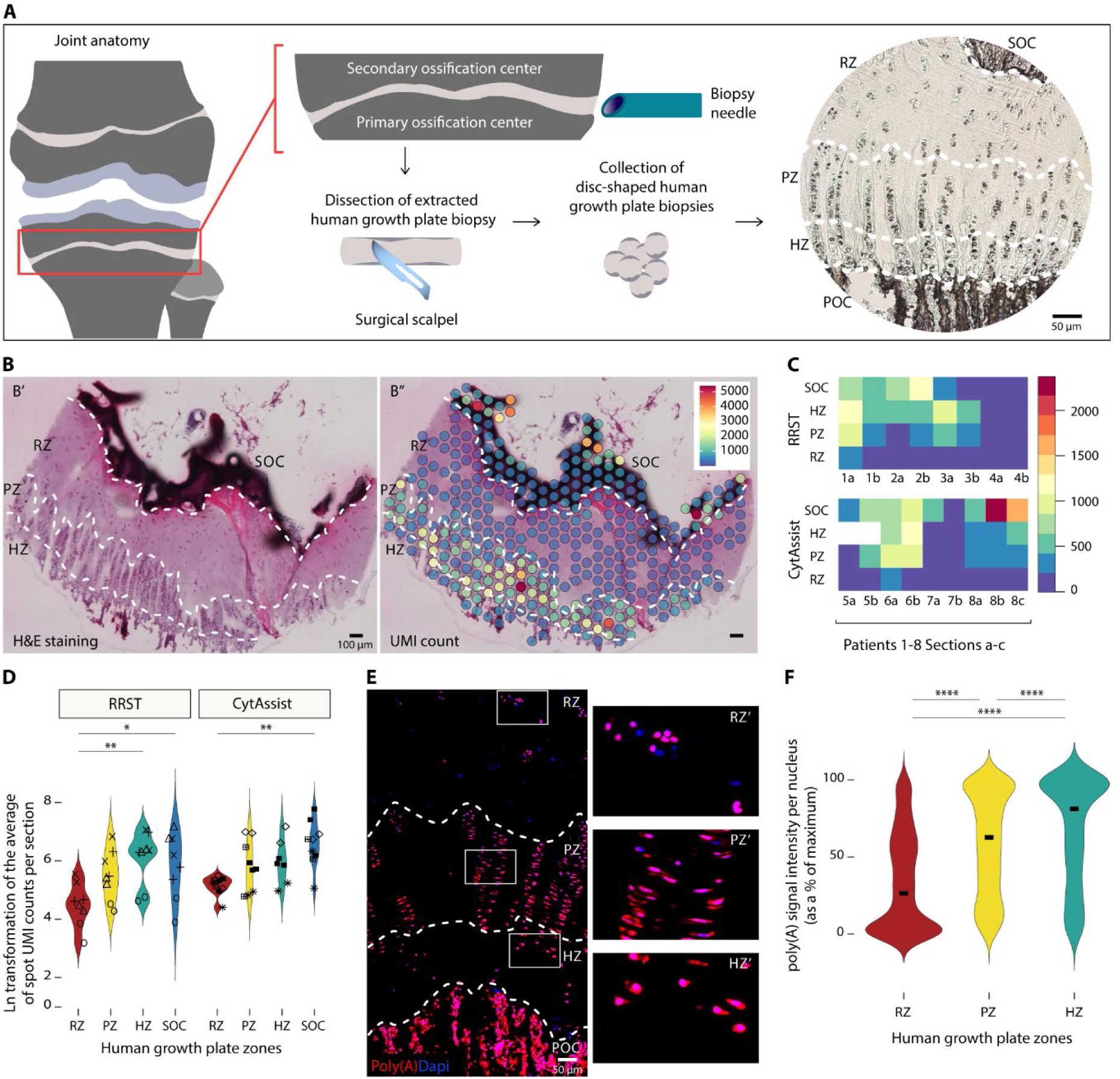
Resting zone chondrocytes contain relatively low amounts of mRNA. (**A**) The procedure for collection of human growth plate biopsies during epiphysiodesis surgeries. (**B**) Hematoxylin and eosin (H&E)-stained tissue section (B’) was histologically separated into areas of growth plate zones and SOC, and spots representing UMI counts from RRST profiling were overlaid (B’’). (**C**) Level-plot of UMI counts per section in patients 1 to 4 for RRST and 5 to 8 for CytAssist. (**D**) Violin plot depicting the Ln transformed average of spot UMI counts in each zone per section in both RRST and CytAssist. The comparison of means between zones in RRST was analyzed using one-way ANOVA (p-value < 0.01) and Tukey post-hoc test: **p < 0.01 and *p < 0.05 from 8 sections from 4 patients. The comparison of means between zones in CytAssist was analyzed using one-way ANOVA (p-value < 0.01) and Tukey post-hoc test: **p < 0.01 from 9 sections from 4 patients (two sections from patient 5 on panel C. shows lack of HZ). (**E**) Representative image of RNAscope assay targeting poly(A) expression in different zones of the human growth plate. (**F**) Mean signal intensity per nucleus was quantified in each zone and converted to a percentage value using the maximum value for each patient. To compare the average signal intensity medians between zones, we used Kruskal-Wallis (p-value < 0.0001) and Wilcoxon post-hoc test: ****p < 0.0001 for all three comparisons. Nuclei from n = 2084 cells in RZ, n = 1985 cells in PZ, n = 784 cells in HZ from 5 patients.

**Table 1.** SRT Exploration dataset. Metadata of the patient information for the tissues analyzed in Figs. 1C, D, 3A-C and Table 2.

| Patient ID | Platform | Genetic sex | Tanner pubertal stage PH<sup>1</sup> | Tanner pubertal stage B<sup>2</sup> | Tanner pubertal stage G<sup>3</sup> | Patient age at surgery |
| --- | --- | --- | --- | --- | --- | --- |
| 1 | RRST | M | 3 | - | 3 | 13 y 2 m |
| 2 | RRST | F | 3 | 4 | - | 12 y 10 m |
| 3 | RRST | M | 3 | - | 3 | 13 y 4 m |
| 4 | RRST | F | 3 to 4 | 4 | - | 12 y 11 m |
| 5 | CytAssist | M | 2 | - | 2 | 12 y 2 m |
| 6 | CytAssist | F | 4 | 4 | - | 14 y 6 m |
| 7 | CytAssist | M | 4 | - | 4 | 13 y 7 m |
| 8 | CytAssist | F | 4 | 4 | - | 14 y 4 m |

*1. Pubic hair. 2. Female breast. 3. Male genital.*

To validate these findings, we utilized the Visium CytAssist Spatial Gene Expression protocol, where an average of three pairs of probes target each transcript (compared to only one pair using RRST), which provided comparable results to RRST experiments. Although the average number of cells captured in each Visium spot was similar throughout the growth plate (fig. S1B), relatively few transcripts were captured in the RZ (Fig. 1B-D, Table 2). To understand the biological relevance of this transcription pattern, we used *in situ* hybridization to visualize mRNA molecules within the human growth plate sections by staining for polyadenylated (poly[A]) RNA tails. This experiment confirmed that RZ cells contained less mRNA than other growth plate chondrocytes (Fig. 1F).

**Table 2.** Average number of UMI counts per spot from SRT. The average of UMI counts per spot was calculated from SRT data from each area of every section. Where sections lacked specific areas, they are denoted NA.

| Platform | RRST |  |  |  |  |  |  |  |  |
| --- | --- | --- | --- | --- | --- | --- | --- | --- | --- |
| PatientID_section | 1_1 | 1_2 | 2_1 | 2_2 | 3_1 | 3_2 | 4_1 | 4_2 |  |
| SOC | 879.51 | 506.78 | 897.15 | 1405.67 | 324.05 | 213.84 | 57.53 | 111.12 |  |
| RZ | 237.90 | 179.59 | 81.47 | 71.07 | 101.13 | 106.97 | 23.81 | 47.35 |  |
| PZ | 893.27 | 369.98 | 157.56 | 190.88 | 551.75 | 237.83 | 68.01 | 83.96 |  |
| HZ | 1078.94 | 505.75 | 512.91 | 525.73 | 1073.84 | 538.24 | 134.29 | 109.61 |  |
| POC | 432.17 | 222 | NA | NA | 1963.05 | 1736.97 | 352.81 | 279.25 |  |
| Platform | CytAssist |  |  |  |  |  |  |  |  |
| PatientID_section | 5_1 | 5_2 | 6_1 | 6_2 | 7_1 | 7_2 | 8_1 | 8_2 | 8_3 |
| SOC | 453.61 | 1007.11 | 790.70 | 1061.44 | 629.73 | 90.81 | 1007.11 | 2652.26 | 1863.61 |
| RZ | 194.33 | 170.33 | 259.01 | 158.69 | 71.97 | 82.98 | 217.63 | 205.93 | 139.76 |
| PZ | 118.80 | 504.45 | 1037.14 | 417.06 | 115.76 | 126.02 | 298.45 | 257.08 | 237.64 |
| HZ | NA | NA | 936.38 | 1429.23 | 188.69 | 148.57 | 299.28 | 422.53 | 633.30 |
| POC | NA | NA | 1281.35 | NA | 428.18 | 285.77 | 1639.79 | 2252.82 | 3721.41 |

To determine whether low RNA expression levels are a feature of RZ cells in another species, we examined RNA content in mouse growth plates (fig. S2A, B). Consistent with our findings in human samples, we observed that mouse RZ cells contained low levels of mRNA molecules in comparison to their neighbors (fig. S2C).

### Resting zone chondrocytes display hallmarks of quiescent stem cells

Having determined via independent methods, and in two species, that RZ cells contain less RNA in comparison to other growth plate chondrocytes, we decided to further investigate the biological relevance of this finding. While quantifying polyadenylated RNA, we noticed that it was predominantly localized in the nucleus of RZ cells in both humans and mice, whereas in other chondrocytes, RNA was distributed between the nucleus and cytoplasm (Fig. 2A, fig. S2B). Upon quantification, the relative amount of RNA found in the nucleus was lower in the PZ and HZ chondrocytes than those in the RZ (Fig. 2A, fig. S2D). The nuclear localization of mRNA is a characteristic of cellular quiescence across multiple species and tissues, including deeply quiescent neural stem cells and hematopoietic stem and progenitor cells (*12*). Quiescence is a state of reversible cell cycle arrest which is a characteristic of long-living adult stem cells, priming them in a “poised” state that enables them to act in tissue homeostasis or regeneration (*13, 14*).

**Figure 2.**
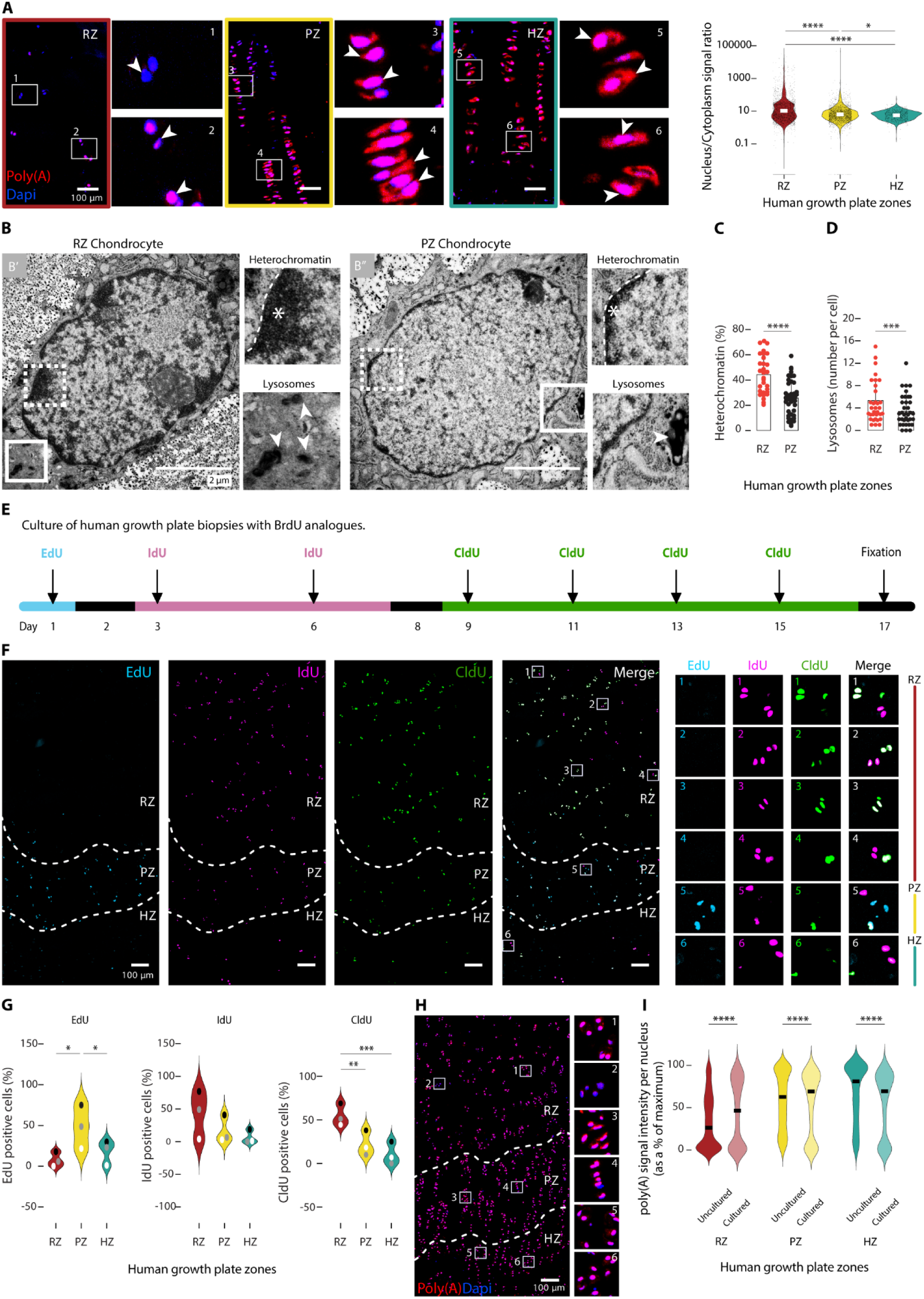
Resting zone chondrocytes display features of cellular quiescence *in vivo*. (**A**) Relative nuclear abundance of poly(A) mRNA was analyzed using Kruskal-Wallis (p-value < 0.0001) and Wilcoxon post-hoc test. The y-axis is plotted on a logarithmic scale with base 10. The median of each zone (RZ = 6.8104, PZ = 5.9712, HZ = 5.7397) is displayed (n = 2860 cells in RZ, n = 2557 cells in PZ, n = 860 cells in HZ from 5 patients). (**B**) TEM images of heterochromatin (asterisks) and lysosomes (arrowheads) in RZ and PZ chondrocytes, from one patient. (**C**) The percentage of each nucleus comprising heterochromatin (n = 31 in RZ, n = 43 in PZ), and (**D**) the number of lysosomes per cell (n = 33 in RZ, n = 44 in PZ) were analyzed using Mann-Whitney test. (**E**) Workflow illustrating the *ex vivo* culture of human growth plate biopsies with thymidine analogues, and (**F**) representative images following visualization. (**G**) Analysis of EdU, IdU and CldU incorporation was conducted using pair-wise matching sample ANOVA (EdU p-value < 0.05, IdU p-value = 0.0989, CldU p-value < 0.001) and Tukey post-hoc test (n = 646 cells in RZ, n = 1227 cells in PZ, n = 505 cells in HZ from 3 patients). (**H**) Poly(A) RNA visualized after culture (**I**) was compared with fresh-frozen slices (presented in Fig. 1F) and analyzed using Mann-Whitney tests. The median of each zone is presented: uncultured RZ = 26.2469, cultured RZ = 46.5278, uncultured PZ = 62.7071, cultured PZ = 69.2647, uncultured HZ = 81.2138, cultured HZ = 69.5188 (for cultured biopsies; n = 1194 cells in RZ, n = 1454 cells in PZ, n = 1053 cells in HZ from 3 patients). In all panels, *p < 0.05, **p < 0.01, ***p < 0.001, ****p < 0.0001.

To further investigate quiescent cells in the growth plate, we conducted transmission electron microscopy (TEM) to compare the ultrastructure of RZ cells with neighboring PZ cells. Firstly, we examined whether human growth plate chondrocytes from the resting and proliferating zones had divergent heterochromatin abundance (Fig. 2B). Notably, RZ chondrocytes had a higher content of normally condensed, electron-dense, heterochromatin (Fig. 2C), which was mainly located at the inner leaflet of the nuclear membrane (Fig. 2B’). In contrast, PZ chondrocytes exhibited discrete areas of heterochromatin within their nuclei (Fig. 2B’’). Consistent with quiescent cells in multiple other tissues (*15, 16*), we found that there were more lysosomes in RZ chondrocytes than PZ chondrocytes (Fig. 2D). Altogether, these results indicate that, as a population, human RZ cells display features indicative of quiescence *in vivo*.

Experiments in mice have revealed that epiphyseal stem cells reside in the RZ (*17, 18*). A subset of slowly-dividing, label-retaining cells (LRCs) in the RZ have been identified by their retention of tritiated thymidine (*19, 20*), BrdU analogues (*17, 21*), or H2B-GFP (*8, 22*) in so-called pulse-chase experiments. While it remains to be determined whether all LRCs are stem cells, progeny of some LRCs can give rise to chondrocyte columns, suggesting that some epiphyseal stem cells are among the LRC (*8, 22*). To determine whether LRCs overlapped with cells containing low levels of mRNA, we combined RNAscope with EdU detection (fig. S3A- C). Some LRCs had elevated mRNA levels, either reflecting that not all LRCs are quiescent, or that they were in the process of leaving quiescence at tissue collection. However, overall, LRCs had significantly lower levels of mRNA than non-LRCs, suggesting that low mRNA levels are a strong indicator of LRCs.

### Human resting zone chondrocytes are functionally quiescent *in vivo*

Relatively low RNA content is an indicator of the G0 phase of the cell cycle in multiple cell types (*23*). To further explore the cell dynamics of human chondrocytes throughout the growth plate, we cultured intact biopsies *ex vivo* in the presence of different BrdU analogues over a time-course (Fig. 2E). We placed freshly collected biopsies directly into EdU-containing medium overnight to capture ongoing cell division events; as anticipated, PZ cells could be labelled using this strategy, whereas few RZ cells incorporated EdU within the first 24 hours of tissue culture (Fig. 2F, G). However, as the culture continued over subsequent days with the inclusion of IdU and CldU in the medium, cells distributed throughout the RZ began to divide (Fig. 2F, G) suggesting that some RZ cells were in a stage of reversible cell cycle arrest *in vivo*. These findings align well with quiescent muscle satellite cells (*24*), hematopoietic stem cells (*25*) and neural stem cells (*26*), which begin dividing when grown *ex vivo*.

To further explore the effects of *ex vivo* culture on the quiescent status of human growth plate chondrocytes, we performed an RNAscope assay targeting poly(A) tails (Fig. 2H). Cultured human biopsies contained a proportion of RZ cells with more mRNA than uncultured biopsies (Fig. 2I), supporting the notion that fewer RZ cells remain quiescent after culture. Although we cannot exclude a prolonged G1 phase in our BrdU-analogue experiments, our combined observations of RNA localization and cellular ultrastructure lead us to propose that a subset of human RZ cells exists in a functionally quiescent state *in vivo*.

### Spatially resolved transcriptomics reveals novel human growth plate markers

Next, we aimed to characterize the transcriptome of the human growth plate and SOC. To this end, we devised a three-step strategy (Fig. 3A) to analyze the SRT data that considered (i) the low number of reads per spot (UMI counts), and (ii) the ability to combine data and results from both platforms (RRST and CytAssist). In the first step, using an “exploration dataset” that contained four different patients per platform (Table 1), we manually annotated the epiphyseal areas (RZ, PZ, HZ, and SOC) in each section to generate a pseudo-count matrix. This matrix sums the reads from all the spots for each gene, area, and sample. This approach is analogous to a pseudo-bulk analysis in single-cell RNA-seq (*27*), which has been shown to reduce false positives in subsequent statistical analyses (*28*). However, in this case the pseudo-bulk was generated by combining the reads from spots instead of cells. Visual inspection of the first two principal components derived from the pseudo-bulk matrix showed that the four distinct areas clustered together as expected (Fig. 3A). In a second step, we identified transcriptomic markers per area by conducting a differential expression analysis between areas. Before investigating the results, we validated this approach by confirming that it could distinguish the zones using well-known cartilage and bone markers (Fig. 3B). As expected, cartilage markers (COL2A1, COL9A1, ACAN, MATN3) were restricted to the growth plate chondrocytes (RZ, PZ plus HZ), and several well-described zone-specific genes were detected in the expected zones (RZ: SFRP5 (*22*), UCMA (*29*); HZ: ALPL (*30*), COL10A1 (*31*)). We also verified the spatial gene expression profiles by generating spatial feature plots for selected markers (Fig. 3C). Altogether, these results demonstrated that SRT generated meaningful results when applied to human growth plate.

**Figure 3.**
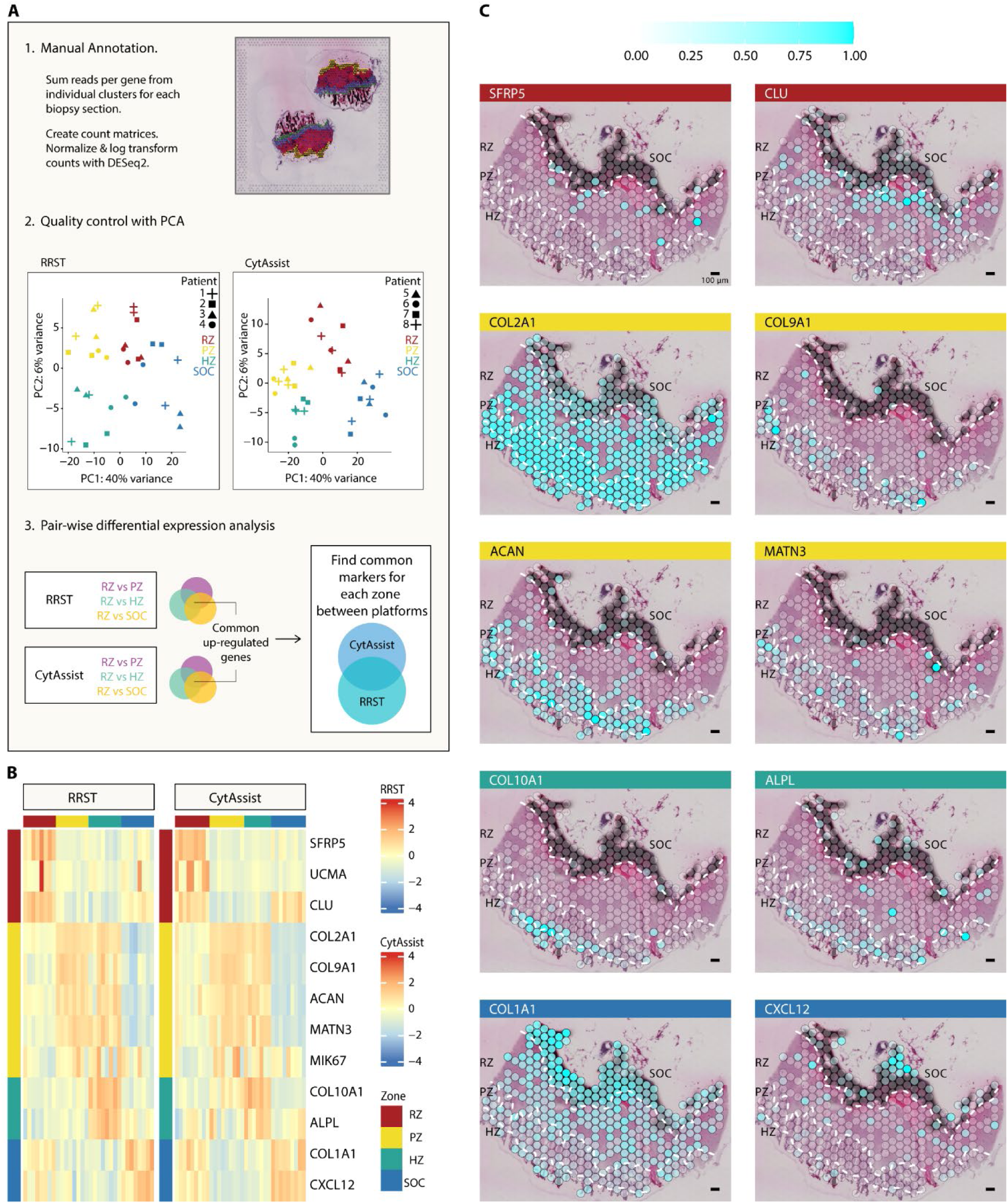
Spatial profiles of cartilage and SOC markers visualized on human growth plate sections. (**A**) Pseudo-Bulk analysis workflow used to analyze SRT data. SRT data collected from RRST and CytAssist platforms was used for transcriptomics analysis. (1) samples were manually annotated based on histology into the different areas: RZ, PZ, HZ, and SOC. Reads per gene from each section were summed per region creating a pseudo-bulk count matrix. Normalization and log transformation of the counts was performed with DESeq2. (2) Principal Component Analysis (PCA) was performed for each platform, where samples grouped distinctly by area. (3) Pair-wise differential expression analysis was performed between the different areas in each platform. Each area-specific marker was significantly up-regulated using both platforms. (**B**) Expression heatmap of 12 selected markers of cartilage and bone in the epiphyseal areas, and (**C**) Spatial feature plot of selected markers of bone and cartilage.

We then applied strict criteria to the pseudo-bulk analysis to explore new markers for the four epiphyseal areas: RZ, PZ, HZ and SOC. In the final step, we considered markers that were consistently profiled and significant across both platforms. As a result, we identified 138 markers that were significantly up-regulated in one area above all others across all eight patients (4 in RZ, 12 in PZ, 28 in HZ and 94 in SOC) (Fig. 4A). We generated spatial feature plots for selected markers to visualize these markers on tissue sections (Fig. 4B), verifying that enrichment of expression occurred predominantly in the identified area.

**Figure 4.**
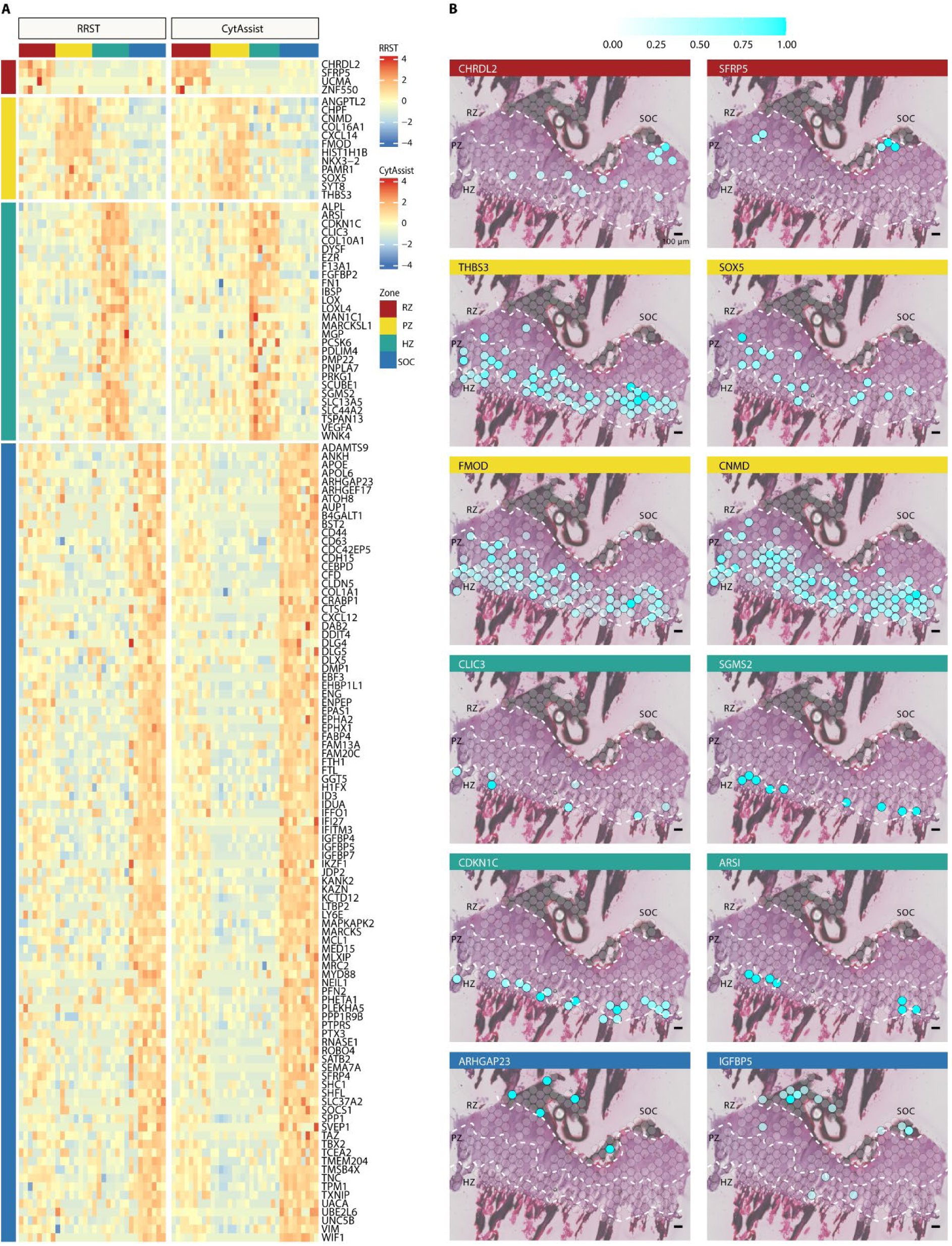
SRT reveals 138 differentially expressed genes between the three growth plate zones and SOC. (**A**) Expression heatmap of 138 genes significantly up-regulated between both platforms in one of the areas above all others. Z-scores for each gene are plotted, blue indicating lower expression and red higher expression of the gene in each sample. Upper bars indicate the sample area labelled, and left bars indicate the area associated for each gene, with RZ in red, PZ in yellow, HZ in green and SOC in blue. (**B**) Spatial feature plot of examples of significantly up-regulated genes for growth plate cartilage and SOC.

We validated these findings in two ways. First, we screened the 138 identified genes in a second cohort (referred to as the “validation dataset” in the materials and methods section, table S1), which included 130 of the markers identified in the exploration dataset (fig. S4A), and which broadly phenocopied the original analysis in the discovery dataset. Secondly, we visualized a marker for each growth plate zone on tissue sections using multi-target RNAscope (fig. S4B). We selected previously unreported markers of the RZ (*CHRDL2*) and HZ (*CLIC3*), as well as one marker we identified in PZ (*THBS3*) that is not a well-known PZ marker but has been previously reported in the murine PZ (*32*) (fig. S4B). These results confirmed that the selected markers were expressed in the cell populations identified by our SRT pseudo-bulk analysis approach, further validating the robustness of our approach and the results.

### Sub-populations of human resting zone chondrocytes have distinct quiescence statuses

To explore the quiescent RZ chondrocytes within the SRT profiles, we used RZ markers to identify sub-populations (fig. S5A) and explored their relative quiescence using spot UMI count. Intriguingly, these markers were able to split the population: spots enriched in CHRDL2 and/or SFRP5 tended to contain relatively high RNA levels, whereas spots enriched in ZNF550 contained relatively low RNA levels (Fig. 5A). Despite applying an analysis approach that identified common markers across all 8 patients, the distributions of these sub-populations between the patients were variable (Fig. 5B), indicating an underlying heterogeneity between individuals, which did not appear to relate to genetic sex or Tanner stage (fig. S5B).

**Figure. 5.**
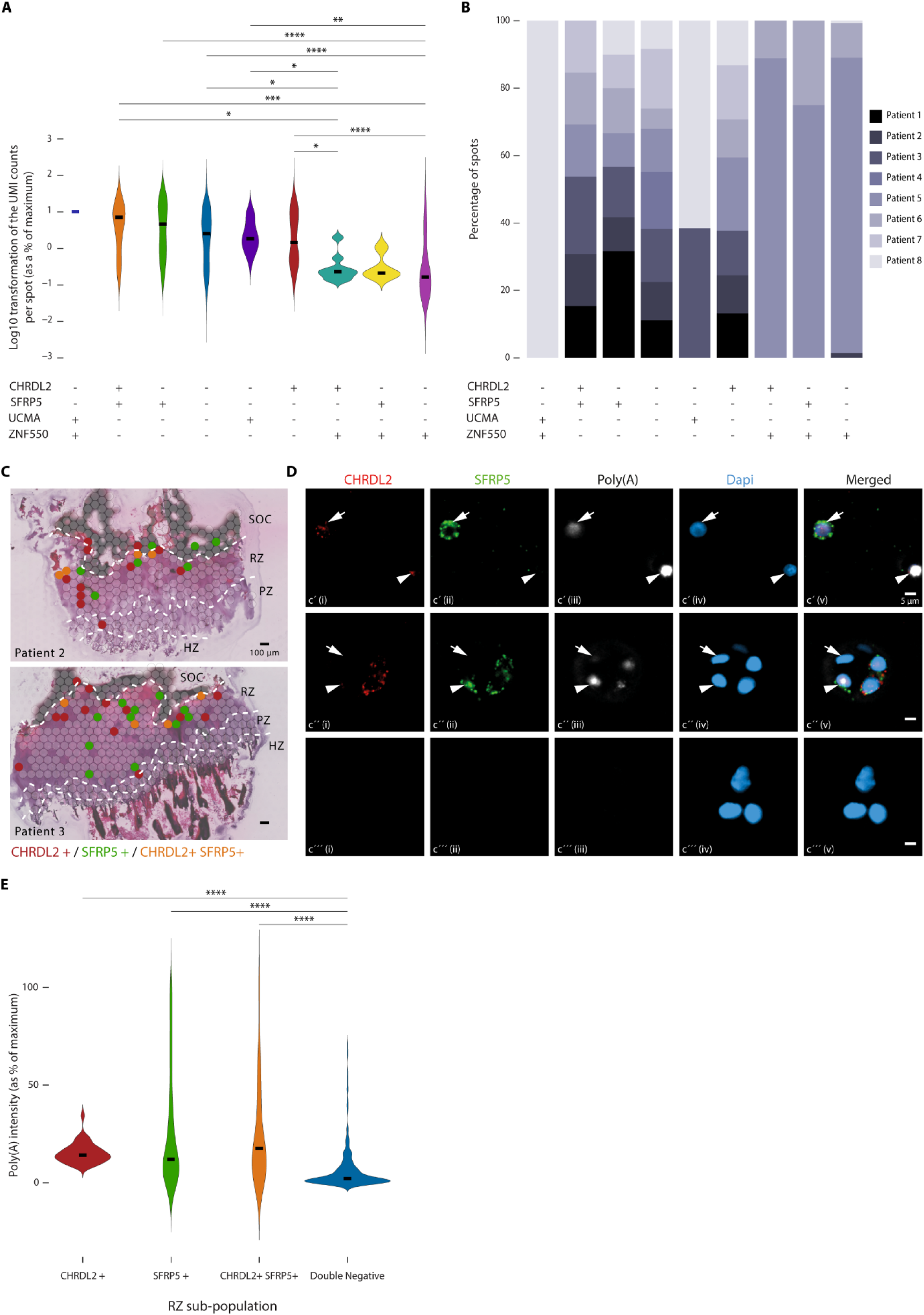
CHRDL2 and SFRP5 can be used to label distinct resting zone sub-populations. (**A**) UMI counts per spot was quantified in the sub-populations of the significantly up-regulated genes in RZ. Medians were statistically compared using Kruskal-Wallis (p-value < 0.0001) and Wilcoxon post-hoc test: *p < 0.05, **p < 0.01, ***p < 0.001, ****p < 0.0001. (**B**) The abundance of the sub-populations in all 8 patients was visualized. (**C**) Spatial feature plot of spot for single expression of CHRDL2 (in red) and SFRP5 (in green) and their co-expression (in orange). Patient 2 is female and patient 3 is male. (**D**) High-magnification images allow visualization of SFRP5/CHRDL2-double-positive cells (arrow d’), -single-positive (arrowheads, d’, d’’) or -double-negative (arrows in d’’ and d’’’). (**E**) Poly(A) signal intensity was quantified in the sub-populations. Medians were statistically compared using Kruskal-Wallis (p-value < 0.001) and Wilcoxon post-hoc test: ****p < 0.0001. The median of each group being CHRDL2+ = 0.1422, SFRP5+ = 0.1200, CHRDL2+ SFRP5+ = 0.1763, Double negative = 0.0210 (n = 381 cells in RZ from 3 patients).

Of the 4 RZ markers identified in SRT data (Fig. 4A), we focused on those directly involved in signal transduction: *SFRP5* (a negative regulator of Wnt signaling), and *CHRDL2* (a bone morphogenic protein [BMP] pathway antagonist), which were identified in 7 of the 8 patients (Fig. 5B). Spots containing either CHRDL2 and/or SFRP5 were spatially distributed throughout the RZ (Fig. 5C). Next, we elaborated on the quiescence statuses of these four sub-populations on the single cell level (Fig. 5D, fig. S5C). RZ cells in which *CHRDL2* and/or *SFRP5* were detected had relatively high mRNA levels (Fig. 5E), confirming that these cells are among the least quiescent RZ cells. Furthermore, RZ chondrocytes within the same cartilage clusters (i.e. chondrons) showed heterogeneity in both marker expression and total mRNA content (Fig. 5D’’). Hence, *SFRP5* and *CHRDL2* can be used to identify sub-populations of human growth plate chondrocytes and were not detected in deeply quiescent RZ cells.

## DISCUSSION

By applying SRT to rare tissues from healthy adolescents, we greatly extended the understanding of the human growth plate transcriptome. We screened the nosology of skeletal dysplasias (*33*) to elaborate on the biological implications of the 138 differentially expressed genes we identified by SRT and found that 14 were included (HZ: *ALPL*, *CDKN1C*, *COL10A1*, *FN1*, *MGP*, *SGMS2*; SOC: *ANKH*, *COL1A1*, *DLX5*, *DMP1*, *FAM20C*, *IDUA*, *LTBP2*, *SFRP4*). All of the listed genes that were enriched in the SOC have previously been shown to have some expression or function in the osteoblast lineage: *ANKH* (*34*) (also reported in chondrocytes and osteoclasts), *COL1A1* (*35*), *DLX5* (*36*) (also reported in chondrocytes (*17*)), *DMP1* (*37*), *FAM20C* (*38*), *IDUA* (*39*), *LTBP2* (*40*), *SFRP4* (*41*), reflecting the presence of mature bone tissue within the SOC.

Within the growth plate, two of the four genes up-regulated in the RZ have been previously detected in corresponding cells in rodents (*Ucma* (*29*), *Sfrp5* (*42*)). The two other genes, *CHRDL2* and *ZNF550*, are newly identified RZ markers, although *Chrdl2* is expressed in RZ cells in mice prior to, but not after SOC formation (*43*). As a BMP pathway antagonist, *CHRDL2* expression in the human RZ fits well with our understanding of BMP signaling in rodents in which BMP antagonists, such as *Bmp3* and *Grem1* (*44*), are expressed in the RZ. These results agree with the overall understanding of BMP signaling in growth plate and shed light on a previously unreported species-specific regulator of human growth.

Several of the genes enriched in the PZ are involved in cartilage formation (e.g. *SOX5* (*45*), *COL16A1* (*46*), *CHPF* (*47*)), a particularly active process in the PZ. Additionally, *Fmod* (*48*) *Cnmd* (*49*) and *Thbs3* (*32*) are enriched in PZ chondrocytes in developing rodents. *NKX3-2* was also significantly enriched in the human PZ. Patients with inactivating mutations in *NKX3-2* develop spondylo-megaepiphyseal-metaphyseal dysplasia (SMMD), which impairs the growth of bones in the neck and trunk whilst those in the appendicular skeleton become longer (*50, 51*). This complicated skeletal phenotype may be because *NKX3-2* is involved in both mesenchymal condensations (*52*) and the repression of chondrocyte hypertrophy (*53*). Although *Nkx3-2* has been detected in PZ chondrocytes in mice limbs (*53*), mouse models of SMMD do not completely phenocopy SMMD patients (*54*). Hence, our results showing that *NKX3-2* is specifically enriched in PZ chondrocytes in adolescent human limbs offers an important perspective of SMMD.

Of the genes enriched in HZ chondrocytes, many have been previously reportedly expressed in the HZ: *ALPL* (*30*), *CDKN1C* (*55*), *COL10A1* (*31*), *FN1* (as well as RZ cells, prior to SOC formation in mice) (*56*), *MGP* (previously reported in PZ and HZ of neonatal mice) (*57*). On the other hand, *WNK4* mutations lead to pseudohypoaldosteronism, involving short stature in a subset of these patients by unknown mechanisms (*58*). We found *WNK4* to be expressed in HZ chondrocytes, which could indicate that *WNK4* has a direct role in short stature via modulation of the growth plate. *SGMS2* is also yet to be described in growth plate cartilage. Mis-sense mutations in *SGMS2*, which encodes the Sphingomyelin Synthase 2, can cause spondylo-metaphyseal dysplasias characterized by short stature, bowed tubular bones and under-mineralization of the skeleton (*59*). To the best of our knowledge, our results are the first to demonstrate *SGMS2* expression in the growth plate, and we have been able to demonstrate its enrichment in hypertrophic chondrocytes of humans. Further research is required to determine if WNK4 and SGMS2 expression/activity in the HZ could account for some of the phenotypes observed in these patients.

To summarize our SRT profiling results, by analyzing data across all 8 patients, we bring to the fore genes likely to be involved in skeletal growth irrespective of genetic sex or Tanner stage. The presence of well-described markers in the anticipated growth plate zones strongly indicates the precision of the SRT methodology and pseudo-bulk analysis approach, strengthening the validity of the less well-described and novel gene expression profiles. Furthermore, these results confirm the relevance of gene expression profiles obtained during decades of animal experimentation to patients. These transcriptional profiles may facilitate a better understanding of diseases, improve the relevance of existing animal studies and facilitate novel strategies to treat these patients.

Based on histological observations, some chondrocytes have been thought of as “resting” since at least the 1920s (*60*). Early experiments using tritiated thymidine to explore cell dynamics revealed the relatively slow rates of division in cells located proximally to the proliferating columns, substantiating their “resting” nature (*19*). RZ chondrocytes are often described by the term “quiescent” in the literature, but whilst they are relatively slowly dividing, other features of cellular quiescence have not been thoroughly explored. Quiescence is an important property of adult stem cells in a variety of tissues, enabling cells to remain in a relatively secure state but be optimally positioned to act in tissue homeostasis or repair (*13, 14*). Our results demonstrate that RZ chondrocytes have the characteristics of quiescent stem cells, including reduced mRNA content (*12*), predominantly nuclear mRNA localization (*12*), increased heterochromatin (*61*), lysosomal accumulation (*15*), and the ability to exit G0 when cultured *ex vivo* (*24, 62*). Hence, our results strengthen the understanding of the quiescent, “resting”, nature of RZ cells and indicate that research demonstrating the presence of skeletal stem cells during postnatal growth in mice, may also be relevant to human skeletal growth.

The murine RZ contains heterogeneous cells expressing a variety of markers including *Sfrp5* (*22*), CD73 (*8, 63*), *Ucma* (*29*), *Axin2* (*63*), *Clu* (*63*), PTHrP (*17, 63*) and *Foxa2* (*18*), among others (*22*). The current consensus is that the RZ chondrocytes include skeletal stem and progenitor cells as the *de facto* stem cells have not yet been identified. Little is known about hierarchy among the populations except for tracing experiments suggesting that *Foxa2*- expressing RZ chondrocytes lie upstream of the PTHrP-positive population (*18*). Our own results demonstrate that, like mice, human RZ chondrocytes are also heterogenous, based on their differential expression of *CHRDL2*, *SFRP5*, *UCMA* and *ZNF550*. To explore the potential hierarchy of these populations, we assessed their quiescence level by UMI count or polyadenylated RNA intensity. Both approaches revealed that RZ sub-populations had heterogenous RNA content, indicating that different sub-populations had varying quiescence statuses. Nevertheless, we determined that sub-populations of RZ cells expressing *CHRDL2* and/or *SFRP5* were located throughout the human RZ and that cells within these sub-populations contained significantly more mRNA than *CHRDL2*/*SFRP5*-double-negative cells. Individual cells located in each chondron were heterogenous for total mRNA, *CHRDL2* and *SFRP5* levels, which could indicate that local regulation within each chondron maintains the RZ, which would align well with our understanding that repressed BMP- and Wnt signaling maintains the RZ in rodents (*22*).

While post-natal growth in mice is fueled by skeletal stem cells (*8*), mouse and human RZ cells have similar characteristics and marker expression (*22*), and it is thought that the general mechanisms of growth between mice and humans are similar (*7*), it is important to emphasize that direct evidence of an epiphyseal stem cell population and niche is lacking. Human growth plates contain relatively large RZs in comparison to rodents, and some of these cells may fulfil roles that have yet to be defined (*9*). Further experiments are required to definitively test if RZ cells, or a specific subset of them, fulfill the necessary criteria to represent *bona fide* stem cells in human growth plates.

### Limitations

Whilst the SRT experiments successfully identified novel growth plate markers, several limitations should be noted. First, the spot size in the SRT we used was 55 µm diameter precluding single cell resolution. Second, due to the relatively low levels of transcription and cell density we needed to devise a robust strategy to detect statistically significant differences between the populations. Furthermore, these SRT methods are probe-based and although covering the whole transcriptome many genes that are important for growth plate biology were missing from one or both panels (e.g. *IHH*, *PTHLH* [encoding PTHrP]). Additionally, although our SRT analysis describes significantly differentially expressed genes across all eight patients, the results were not sufficient to determine those specific to, for example, genetic sex or pubertal stage, as determined by our power analyses.

In summary, we greatly elaborated upon the human growth plate transcriptome, allowing us to identify novel zone-specific markers, new primary growth disorders, and candidate pharmacological targets. Furthermore, our results revealed that the RZ contains chondrocytes that display characteristics of adult stem cells and sub-populations of varying quiescence status. Together, these findings can facilitate improved diagnosis and treatment strategies of patients with skeletal growth disorders.

## MATERIALS AND METHODS

### Study Design

In this study, we aimed at deciphering the transcriptional profiles of the human growth plate. We selected biopsies containing growth plate and SOC for SRT. We applied RRST, a Visium method we previously optimized for use with cartilage tissue, and to improve sequencing results, complemented this with CytAssist. To analyze samples from both RRST and CytAssist we selected the 4 patients from each platform with highest median UMI count per spot, resulting in an exploration dataset of 4 male and 4 female patients, 2 per platform. All other sequenced sections using RRST were added to the Validation cohort. Spots were manually assigned into zones by a trained observer based. No statistical method was used to pre-determine sample size. The experiments were not randomized, and the investigators were not blinded to allocation during experiments and outcome assessment. Animal experiments were performed in accordance with the ARRIVE guidelines (*64*).

### Ethics declaration

The collection of human growth plate samples was approved by the local human ethics committee (KI forskningsetikkommitté Nord vid Karolinska Sjukhuset), ethical permit 97/214. Informed consent was obtained from all participating subjects and their parents before enrolment in the study, which was performed according to the Declaration of Helsinki. Mouse bone samples were collected according to ethical permit 16673/2020, approved by Stockholm’s animal experiment ethics committee (Stockholms djurförsöksetiska nämnd).

### Human growth plate sample information and collection

Human growth plate samples were collected from constitutionally tall-stature patients undergoing epiphyseal surgery (epiphysiodesis) to reduce bone growth at the Karolinska University Hospital. To be eligible for treatment a predicted adult height of at least 186 cm for girls or 200 cm for boys (corresponding to 3 standard deviations above mean height), at least 8 cm of remaining predicted growth, and a strong desire to undergo surgery must exist. Immediately after collection, the biopsies from distal femora and proximal tibiae were transferred into DMEM-F12 medium (phenol red free [21041025; Gibco]) on ice and transported directly to the laboratory. Under microscopy, biopsies in DMEM/F12 were dissected on ice using a scalpel, into slices of approximately 1 mm thickness. For SRT experiments, fresh-frozen samples were prepared by placing each slice into Optimal Cutting Temperature (OCT) compound medium (4583; Tissue-Tek) within a cryomold (62534-10; Tissue-Tek), which was rapidly frozen in a hexane (296090-1; Sigma-Aldrich) bath that had been pre-cooled in a tank of absolute ethanol (01399; Histolab) containing dry-ice (carbon dioxide ice). For RNAscope, each slice was fixed in pre-cooled 4 % formaldehyde (64833; Merck)/PBS for 48 hours at 4 °C. Samples were then placed into 30 % sucrose (S-2885; AG Scientific) overnight and embedded in OCT medium in cryomolds. Samples were stored at - 80°C prior to cryo-sectioning.

### RRST Gene Expression library preparation

Fresh-frozen human growth plate slices were cryo-sectioned in a CryoStar NX70 (957070; Epredia) at 10 µm thickness, placed onto glass slides from Visium Spatial for FFPE Gene Expression kit (PN-1000338; 10x Genomics), and stored at −80°C before processing. Each Capture Area contained section(s) from 1 patient. Spatial gene expression libraries were generated following RRST Gene Expression protocol, as previously reported(*11*). Final libraries were purified and assessed using an Agilent Bioanalyzer (Agilent 2100 Bioanalyzer system) and quantified using the Qubit Fluorometric quantification (ThermoFisher) before sequencing on NextSeq2000 (Illumina) at a depth of 25,000 reads per tissue covered spot on Capture Area. The Read 1 (28 bp [base pairs]) encodes for the Spatial barcode and Unique Molecular Identifier (UMI) and Read 2 (50 bp) was used to sequence the ligated probe insert.

### CytAssist Gene Expression library preparation

Fresh-frozen human growth plate slices were cryo-sectioned at 10 µm thickness, placed within a 6.5 mm Capture Area on SuperFrost Ultra Plus GOLD Adhesion slides (11976299; Thermo Fisher Scientific) following 10x Genomics Tissue Preparation Guide (Demonstrated Protocol, CG000518 Rev C), and stored in −80°C freezer. Each Capture Area contained sections from two patients. The tissue slides were removed from −80 °C freezer, fixed and stained as in RRST protocol, as previously reported (*11*). CytAssist gene expression libraries were generated using Visium CytAssist Spatial Gene Expression for FFPE kit (PN-1000520; 10x Genomics) following 10x Genomics Visium Spatial Gene expression protocol (User Guide, CG000495 Rev D). Final libraries were purified and assessed using an Agilent Bioanalyzer (Agilent 2100 Bioanalyzer system) and quantified using the Qubit Fluorometric Quantification (ThermoFisher) before sequencing on NextSeq2000 (Illumina) at a depth of 25,000 reads per tissue covered spot on Capture Area. The Read 1 (28 bp) encodes for the Spatial barcode and UMI and Read 2 (50 bp) was used to sequence the ligated probe insert.

### Data pre-processing

Sequenced libraries were processed using Space Ranger software (version 1.3.1 for RRST data and version 2.0.1 for CytAssist data, 10x Genomics). Reads were aligned to the pre-built human reference genome the GRCh38, provided by 10x Genomics (version refdata-gex-GRCh38-2020-A). h5 files and image data were obtained.

### Spot-level annotation

Spots from all Capture Areas were annotated on a spot-by-spot basis using Loupe Browser version 6.2.0 (10x Genomics). Based on tissue histology, spots were manually defined into areas: SOC, RZ, PZ, HZ, and POC. Spots lacking cells were excluded. Since human growth plate slices did not always contain entire growth plates, we prioritized the use of sections that contained the complete SOC and RZ with SRT. Of those used to generate the exploration dataset, 29.4 % of tissue sections lacked POC (Table 2), which was excluded from downstream analyses.

### Bioinformatics analysis

The h5 files obtained from Space Ranger were converted into Seurat objects. The table containing the area annotation was added to the metadata for each sample and section. Finally, Seurat objects were merged to obtain a Seurat object containing all the samples. For each section, the raw reads coming from spots labelled as SOC, RZ, PZ and HZ were summed up generating a new count matrix (pseudo-bulk approach). RRST and CytAssist samples were split into two count matrices and a metadata table was generated for each platform. Analyses for each sample were done independently following the same pipeline. First, only genes with at least 10 reads among all the samples were kept for further analysis per platform. Patient-based batch correction was performed using ComBat_seq (*65*). Normalization and differential expression analysis were performed using DESeq2 separately for each platform (*66*). Pairwise comparisons between the different areas were performed and for each pairwise comparison genes with a *p*-value < 0.2 and that were either up- or down-regulated in the analysis associated with both platforms were combined using the fisher test (*67*).

A gene was considered a candidate marker of an area (SOC, RZ, PZ or HZ) if it was up-regulated in the specific area compared to the other three areas and has a False Discovery Rate (FDR) adjusted Fisher *p* value < 0.05 in the three comparisons; based on the Benjamini Hochberg threshold. For validation purposes, the same pseudo-bulk strategy was followed in the validation dataset. In this case we only checked the differential expression of the 138 markers genes found in the exploration dataset. These samples were all sequenced on RRST platform; similar normalization and differential analysis were conducted. Genes were considered significant with an FDR < 0.05. Only 130 of the 138 genes are presented in Fig. S4A since the other 8 genes did not meet the expression level threshold in the validation dataset.

### RNAscope *in situ* hybridization

Cryo-sections at 10 μm thickness were prepared from blocks containing 48-hour fixed samples using Leica CM3050 S Cryostat (LeicaBiosystems), placed onto SuperFrost Ultra Plus GOLD Adhesion slides (11976299; Thermo Fisher Scientific), and stored at −80°C before processing. To visualize RNA molecules on tissue section we utilized RNAscope® as previously described (*68*), outlined in the supplementary methods section. The RNA expression levels for our target were investigated using RNAscope® Multiplex Fluorescent Detection Reagents v2 kit (323100; ACD) and RNAscope® 4-Plex Ancillary kit (323120; ACD) using probes for poly(A) (423731; ACD), hh-CHRDL2 (1043521-C1; ACD), hh-THBS3 (1274141-C2; ACD), hh-CLIC3 (569631-C3; ACD), and hh-SFRP5 (1056891-C4; ACD). Positive RNA control probe (320881; ACD) and negative RNA control probe (320871; ACD) were used to validate the RNAscope assay on mouse or human sections.

### Triple labelling of human growth plate biopsies with thymidine analogues

Individual human growth plate slices were grown in one well of a 24-well plate (83.3922; Sarstedt) with 2 ml of culture medium containing DMEM/F12 (11320033; Gibco) supplemented with 50 µg/ml ascorbic acid (A9560; Sigma-Aldrich), 0.2 % BSA (05470-5G; Sigma-Aldrich) and 50 µg/ml gentamicin (11530506; Fisher Scientific). EdU was added on day 1 at 1:1000 of 25 mg/mL stock solution and incubated overnight. IdU (I7125-5Gl; Merck) was added on days 3 and 6 at 1:1000 of 250 mM stock solution. CldU (C6891; Merck) was added on days 9, 11, 13, and 15 at 1:1000 of 15 mM stock solution. On days 2 and 8, wells were rinsed with PBS and slices were placed in culture medium without thymidine analogues. Explants were maintained in a humidified atmosphere (37°C, 5% CO_2_) within an incubator (51033595; Thermo Scientific). Explants were typically placed in culture within approximately two hours of surgery. On day 17, the explants were briefly rinsed in PBS and fixed in pre-cooled 4 % PFA/PBS overnight at 4 °C. Subsequently, the samples were incubated in 30 % sucrose overnight at 4 °C, and the following day they were embedded in OCT in cryomolds. Samples were stored at −80 °C before cryo-sectioning.

The detection method was used as described previously (*8*), with modifications made because some fluorescent antibodies were no longer commercially available. CldU was visualized using anti-BrdU antibodies conjugated to Alexa Fluor 488 ([clone BU1/75 (ICR1)] 220074l; Abcam). The anti-IdU antibodies were detected using donkey anti-mouse antibodies conjugated to Alexa Fluor 555 (150106; Abcam), and EdU was detected using Alexa Fluor azide 647.

### Statistical analysis

Normality was assessed using the Shapiro-Wilk test; where normality could be assumed, parametric tests were performed (Figs. 1D, 2G), otherwise non-parametric tests (Figs. 1F, 2A, 2C, 2D, 2I, 4D, S1B, S2C, S2D, S3C) were applied. Data were processed using RStudio software version 4.1.2 (2021-11-01) or GraphPad Prism version 10.2.3.

## ACKNOWLEDGEMENTS

We thank Dr. Henrik Wehtye for methodological support and Dr. Per Roos for translation assistance. We would like to thank National Genomics Infrastructure (NGI), Sweden, for providing infrastructure support. We also acknowledge support of the Bioimaging core facility and the electron microscopy unit Emil at Karolinska Institutet.

## AUTHOR CONTRIBUTIONS

<u>Conceptualization</u> Lead: PTN; supporting: JL, RM, DG-C; <u>Methodology</u> lead: MA, LAG; supporting: ARLP, LSud, JGM-A, ŽA, YZ, FZ, RM, DG-C, PTN; <u>Software</u> lead: DG-C, ARLP; supporting: LSud, LL, MA; <u>Validation</u> lead: MA, ARLP; supporting: LAG; <u>Formal</u> <u>analysis</u> lead: MA, ARLP, LSud; supporting: JGM-A, LL, DG-C, PTN; <u>Investigation</u> lead: MA, ARLP, LAG, RM, PTN, DG-C; supporting: ŽA, LL, HQ, KB, JNT, LSäv, FP ; <u>Resources</u> lead: JL, LSäv; supporting: PTN, FP, JNT, DG-C; <u>Data Curation</u> lead: MA, ARLP, LAG, LS; supporting: JGM-A; <u>Writing - Original Draft</u> lead: PTN; supporting: MA, ARLP, DG-C; <u>Writing - Review & Editing</u> lead: PTN, MA, DG-C; supporting: RM, HQ, ARLP, LSäv, JGM- A; <u>Visualization</u> lead: MA, ARLP; supporting: JGM-A, AN; <u>Supervision</u> lead: PTN, DG-C; supporting: KB, HQ, JL, RM; <u>Project administration</u> lead: PTN, DG-C, RM; supporting: KB, HQ, JL; <u>Funding acquisition</u> lead: PTN, JL, DG-C

## REFERENCES

1. H. M. Kronenberg, Developmental regulation of the growth plate. Nature 423, 332–336 (2003).

2. A. S. Chagin, P. T. Newton, Postnatal skeletal growth is driven by the epiphyseal stem cell niche: potential implications to pediatrics. Pediatr Res 87, 986–990 (2020).

3. G.L. Galea, M.R. Zein, S. Allen, P. Francis-West, Making and shaping endochondral and intramembranous bones. Dev Dyn. 250 (3):414–449 (2021).

4. M. Haymond, A.-M. Kappelgaard, P. Czernichow, B. M. K. Biller, K. Takano, W. Kiess, T. participants in the global advisory panel meeting on the effects of growth hormone, Early recognition of growth abnormalities permitting early intervention. Acta Paediatr 102, 787– 796 (2013).

5. J. M. Wit, S. D. Joustra, Long-acting PEGylated growth hormone in children with idiopathic short stature: time to reconsider our diagnostic and treatment policy? Eur J Endocrinol 188, R1–R4 (2023).

6. E. Benyi, M. Berner, I. Bjernekull, A. Boman, D. Chrysis, O. Nilsson, A. Waehre, H. Wehtje, L. Sävendahl, Efficacy and Safety of Percutaneous Epiphysiodesis Operation around the Knee to Reduce Adult Height in Extremely Tall Adolescent Girls and Boys. Int J Pediatr Endocrinol 2010, 740629 (2010).

7. M. Xie, P. Gol’din, A. N. Herdina, J. Estefa, E. V Medvedeva, L. Li, P. T. Newton, S. Kotova, B. Shavkuta, A. Saxena, L. T. Shumate, B. D. Metscher, K. Großschmidt, S. Nishimori, A. Akovantseva, A. P. Usanova, A. D. Kurenkova, A. Kumar, I. L. Arregui, P. Tafforeau, K. Fried, M. Carlström, A. Simon, C. Gasser, H. M. Kronenberg, M. Bastepe, K. L. Cooper, P. Timashev, S. Sanchez, I. Adameyko, A. Eriksson, A. S. Chagin, Secondary ossification center induces and protects growth plate structure. Elife 9:e55212 (2020).

8. P. T. Newton, L. Li, B. Zhou, C. Schweingruber, M. Hovorakova, M. Xie, X. Sun, L. Sandhow, A. V Artemov, E. Ivashkin, S. Suter, V. Dyachuk, M. El Shahawy, A. Gritli-Linde, T. Bouderlique, J. Petersen, A. Mollbrink, J. Lundeberg, G. Enikolopov, H. Qian, K. Fried, M. Kasper, E. Hedlund, I. Adameyko, L. Sävendahl, A. S. Chagin, A radical switch in clonality reveals a stem cell niche in the epiphyseal growth plate. Nature 567, 234–238 (2019).

9. N. F. Kember, H. A. Sissons, Quantitative histology of the human growth plate. Journal of Bone and Joint Surgery - Series B 58 (1976).

10. B. C. J. van der Eerden, M. Karperien, J. M. Wit, Systemic and Local Regulation of the Growth Plate. Endocr Rev 24, 782–801 (2003).

11. R. Mirzazadeh, Z. Andrusivova, L. Larsson, P. T. Newton, L. A. Galicia, X. M. Abalo, M. Avijgan, L. Kvastad, A. Denadai-Souza, N. Stakenborg, A. B. Firsova, A. Shamikh, A. Jurek, N. Schultz, M. Nistér, C. Samakovlis, G. Boeckxstaens, J. Lundeberg, Spatially resolved transcriptomic profiling of degraded and challenging fresh frozen samples. Nat Commun 14, 509 (2023).

12. A. Rossi, A. Coum, M. Madelénat, L. Harris, A. Miedzik, S. Strohbuecker, A. Chai, H. Fiaz, R. Chaouni, P. Faull, W. Grey, D. Bonnet, F. Hamid, E. Makeyev, A. Snijders, G. Kelly, F. Guillemot, R. Sousa-Nunes, Neural stem cells alter nucleocytoplasmic partitioning and accumulate nuclear polyadenylated transcripts during quiescence. bioRxiv (2021), doi:10.1101/2021.01.06.425462.

13. C. T. J. van Velthoven, T. A. Rando, Stem Cell Quiescence: Dynamism, Restraint, and Cellular Idling. Cell Stem Cell 24 (2):213–225 (2019).

14. A. de Morree, T. A. Rando, Regulation of adult stem cell quiescence and its functions in the maintenance of tissue integrity. Nat Rev Mol Cell Biol 24, 334–354 (2023).

15. D. S. Leeman, K. Hebestreit, T. Ruetz, A. E. Webb, A. McKay, E. A. Pollina, B. W. Dulken, X. Zhao, R. W. Yeo, T. T. Ho, S. Mahmoudi, K. Devarajan, E. Passegué, T. A. Rando, J. Frydman, A. Brunet, Lysosome activation clears aggregates and enhances quiescent neural stem cell activation during aging. Science (1979) 359 (2018).

16. C. Settembre, R. M. Perera, Lysosomes as coordinators of cellular catabolism, metabolic signalling and organ physiology. Nat Rev Mol Cell Biol 25, 223–245 (2024).

17. K. Mizuhashi, W. Ono, Y. Matsushita, N. Sakagami, A. Takahashi, T. L. Saunders, T. Nagasawa, H. M. Kronenberg, N. Ono, Resting zone of the growth plate houses a unique class of skeletal stem cells. Nature 563, 254–258 (2018).

18. S. Muruganandan, R. Pierce, D. A. Teguh, R. F. Perez, N. Bell, B. Nguyen, K. Hohl, B. D. Snyder, M. W. Grinstaff, H. Alberico, D. Woods, Y. Kong, C. Sima, S. Bhagat, K. Ho, V. Rosen, L. Gamer, A. M. Ionescu, A FoxA2+ long-term stem cell population is necessary for growth plate cartilage regeneration after injury. Nat Commun 13, 2515 (2022).

19. N. F. Kember, Cell division in endochondral ossification. Journal of Bone & Joint Surgery, British Volume 42-B, 824–839 (1960).

20. N. F. Kember, B. E. Lambert, Slowly Cycling Cells In Growing Bone. Cell Prolif 14, 327–330 (1981).

21. A. S. Chagin, K. K. Vuppalapati, T. Kobayashi, J. Guo, T. Hirai, M. Chen, S. Offermanns, L. S. Weinstein, H. M. Kronenberg, G-protein stimulatory subunit alpha and Gq/11α G-proteins are both required to maintain quiescent stem-like chondrocytes. Nat Commun 5, 1–14 (2014).

22. S. A. Hallett, Y. Matsushita, W. Ono, N. Sakagami, K. Mizuhashi, N. Tokavanich, M. Nagata, A. Zhou, T. Hirai, H. M. Kronenberg, N. Ono, Chondrocytes in the resting zone of the growth plate are maintained in a wnt-inhibitory environment. Elife 10 (2021), doi:10.7554/eLife.64513.

23. K. Toba, E. F. Winton, T. Koike, A. Shibata, Simultaneous three-color analysis of the surface phenotype and DNA-RNA quantitation using 7-amino-actinomycin D and pyronin Y. J Immunol Methods 182 (2):193–207 (1995).

24. M. Quarta, J. O. Brett, R. DiMarco, A. De Morree, S. C. Boutet, R. Chacon, M. C. Gibbons, V. A. Garcia, J. Su, J. B. Shrager, S. Heilshorn, T. A. Rando, An artificial niche preserves the quiescence of muscle stem cells and enhances their therapeutic efficacy. Nat Biotechnol 34, 752–759 (2016).

25. H. Kobayashi, T. Morikawa, A. Okinaga, F. Hamano, T. Hashidate-Yoshida, S. Watanuki, D. Hishikawa, H. Shindou, F. Arai, Y. Kabe, M. Suematsu, T. Shimizu, K. Takubo, Environmental Optimization Enables Maintenance of Quiescent Hematopoietic Stem Cells Ex Vivo. Cell Rep 28, 145–158.e9 (2019).

26. E. Csaszar, D. C. Kirouac, M. Yu, W. Wang, W. Qiao, M. P. Cooke, A. E. Boitano, C. Ito, P. W. Zandstra, Rapid Expansion of Human Hematopoietic Stem Cells by Automated Control of Inhibitory Feedback Signaling. Cell Stem Cell 10, 218–229 (2012).

27. A. T. L. Lun, J. C. Marioni, Overcoming confounding plate effects in differential expression analyses of single-cell RNA-seq data. Biostatistics 18 (3):451–464 (2017).

28. J. W. Squair, M. Gautier, C. Kathe, M. A. Anderson, N. D. James, T. H. Hutson, R. Hudelle, T. Qaiser, K. J. E. Matson, Q. Barraud, A. J. Levine, G. La Manno, M. A. Skinnider, G. Courtine, Confronting false discoveries in single-cell differential expression. Nat Commun 12, 5692 (2021).

29. A. Tagariello, J. Luther, M. Streiter, L. Didt-Koziel, M. Wuelling, C. Surmann-Schmitt, M. Stock, N. Adam, A. Vortkamp, A. Winterpacht, Ucma - A novel secreted factor represents a highly specific marker for distal chondrocytes. Matrix Biology 27, 3–11 (2008).

30. D. Miao, A. Scutt, Histochemical localization of alkaline phosphatase activity in decalcified bone and cartilage. Journal of Histochemistry and Cytochemistry 50 (2002).

31. J. Chen, F. Chen, H. Bian, Q. Wang, X. Zhang, L. Sun, J. Gu, Y. Lu, Q. Zheng, Hypertrophic chondrocyte-specific Col10a1 controlling elements in Cre recombinase transgenic studies. Am J Transl Res 11 (10):6672–6679 (2019).

32. M. L. Iruela-Arispe, D. A. J. Liska, E. H. Sage, P. Bornstein, Differential expression of thrombospondin 1, 2, and 3 during murine development. Developmental Dynamics 197 (1):40–56 (1993).

33. M. L. Warman, V. Cormier-Daire, C. Hall, D. Krakow, R. Lachman, M. LeMerrer, G. Mortier, S. Mundlos, G. Nishimura, D. L. Rimoin, S. Robertson, R. Savarirayan, D. Sillence, J. Spranger, S. Unger, B. Zabel, A. Superti-Furga, Nosology and classification of genetic skeletal disorders: 2010 revision. Am J Med Genet A 155, 943–968 (2011).

34. I. P. Chen, L. Wang, X. Jiang, H. L. Aguila, E. J. Reichenberger, A Phe377del mutation in ANK leads to impaired osteoblastogenesis and osteoclastogenesis in a mouse model for craniometaphyseal dysplasia (CMD). Hum Mol Genet 20 (5):948–61 (2011).

35. I. Kalajzic, Z. Kalajzic, M. Kaliterna, G. Gronowicz, S. H. Clark, A. C. Lichtler, D. Rowe, Use of type I collagen green fluorescent protein transgenes to identify subpopulations of cells at different stages of the osteoblast lineage. Journal of Bone and Mineral Research 17 (1):15–25 (2002).

36. N. Samee, V. Geoffroy, C. Marty, C. Schiltz, M. Vieux-Rochas, G. Levi, M. C. De Vernejoul, Dlx5, a positive regulator of osteoblastogenesis, is essential for osteoblast-osteoclast coupling. American Journal of Pathology 173 (3):773–80 (2008).

37. I. Kalajzic, A. Braut, D. Guo, X. Jiang, M. S. Kronenberg, M. Mina, M. A. Harris, S. E. Harris, D. W. Rowe, Dentin matrix protein 1 expression during osteoblastic differentiation, generation of an osteocyte GFP-transgene. Bone 35 (1):74–82 (2004).

38. K. Hirose, T. Ishimoto, Y. Usami, S. Sato, K. Oya, T. Nakano, T. Komori, S. Toyosawa, Overexpression of Fam20C in osteoblast in vivo leads to increased cortical bone formation and osteoclastic bone resorption. Bone 138, 115414 (2020).

39. S. C. Kuehn, T. Koehne, K. Cornils, S. Markmann, C. Riedel, J. M. Pestka, M. Schweizer, C. Baldauf, T. A. Yorgan, M. Krause, J. Keller, M. Neven, S. Breyer, R. Stuecker, N. Muschol, B. Busse, T. Braulke, B. Fehse, M. Amling, T. Schinke, Impaired bone remodeling and its correction by combination therapy in a mouse model of mucopolysaccharidosis-I. Hum Mol Genet 24 (24):7075–86 (2015).

40. J. R. Schmidt, S. Kliemt, C. Preissler, S. Moeller, M. Von Bergen, U. Hempel, S. Kalkhof, Osteoblast-released matrix vesicles, regulation of activity and composition by sulfated and non-sulfated glycosaminoglycans. Molecular and Cellular Proteomics 15 (2):558–72 (2016).

41. G. J. Spencer, J. C. Utting, S. L. Etheridge, T. R. Arnett, P. G. Genever, Wnt signalling in osteoblasts regulates expression of the receptor activator of NFκB ligand and inhibits osteoclastogenesis in vitro. J Cell Sci 119 (7):1283–1296 (2006).

42. J. C. K. Lui, A. C. Andrade, P. Forcinito, A. Hegde, W. P. Chen, J. Baron, O. Nilsson, Spatial and temporal regulation of gene expression in the mammalian growth plate. Bone 46 (5): 1380–1390 (2010).

43. N. Nakayama, C. Y. E. Han, L. Cam, J. I. Lee, J. Pretorius, S. Fisher, R. Rosenfeld, S. Scully, R. Nishinakamura, D. Duryea, G. Van, B. Bolon, T. Yokota, K. Zhang, A novel chordin-like BMP inhibitor, CHL2, expressed preferentially in chondrocytes of developing cartilage and osteoarthritic joint cartilage. Development 131 (1): 229–240 (2004).

44. P. Garrison, S. Yue, J. Hanson, J. Baron, J. C. Lui, Spatial regulation of bone morphogenetic proteins (BMPs) in postnatal articular and growth plate cartilage. PLoS One 12 (5):e0176752 (2017).

45. P. Smits, P. Li, J. Mandel, Z. Zhang, J. M. Deng, R. R. Behringer, B. de Crombrugghe, V. Lefebvre, The Transcription Factors L-Sox5 and Sox6 Are Essential for Cartilage Formation. Dev Cell 1, 277–290 (2001).

46. A. Kassner, U. Hansen, N. Miosge, D. P. Reinhardt, T. Aigner, L. Bruckner-Tuderman, P. Bruckner, S. Grässel, Discrete integration of collagen XVI into tissue-specific collagen fibrils or beaded microfibrils. Matrix Biology 22 (2): 131–143 (2003).

47. H. Ogawa, S. Hatano, N. Sugiura, N. Nagai, T. Sato, K. Shimizu, K. Kimata, H. Narimatsu, H. Watanabe, Chondroitin Sulfate Synthase-2 Is Necessary for Chain Extension of Chondroitin Sulfate but Not Critical for Skeletal Development. PLoS One 7 (8):e43806 (2012).

48. C. Itzstein, H. Cheynel, B. Burt-Pichat, B. Merle, L. Espinosa, P. D. Delmas, C. Chenu, Fibromodulin is expressed by both chondrocytes and osteoblasts during fetal bone development. J Cell Biochem 82 (1):46–57 (2001).

49. B. Amil, M. Fernandez-Fuente, I. Molinos, J. Rodriguez, E. Carbajo-Pérez, E. Garcia, T. Yamamoto, F. Santos, Chondromodulin-I expression in the growth plate of young uremic rats. Kidney Int 66, 51–59 (2004).

50. J. Hellemans, M. Simon, A. Dheedene, Y. Alanay, E. Mihci, L. Rifai, A. Sefiani, Y. van Bever, M. Meradji, A. Superti-Furga, G. Mortier, Homozygous Inactivating Mutations in the NKX3-2 Gene Result in Spondylo-Megaepiphyseal-Metaphyseal Dysplasia. The American Journal of Human Genetics 85, 916–922 (2009).

51. P. O. Simsek-Kiper, C. Kosukcu, O. Akgun-Dogan, R. Gocmen, G. E. Utine, T. Soyer, A. Korkmaz-Toygar, G. Nishimura, M. Alikasifoglu, K. Boduroglu, A novel NKX3-2 mutation associated with perinatal lethal phenotype of spondylo-megaepiphyseal-metaphyseal dysplasia in a neonate. Eur J Med Genet 62, 21–26 (2019).

52. C. Tribioli, M. Frasch, T. Lufkin, Bapxl: an evolutionary conserved homologue of the Drosophila bagpipe homeobox gene is expressed in splanchnic mesoderm and the embryonic skeleton. Mech Dev 65, 145–162 (1997).

53. S. Provot, H. Kempf, L. C. Murtaugh, U. Chung, D.-W. Kim, J. Chyung, H. M. Kronenberg, A. B. Lassar, Nkx3.2/Bapx1 acts as a negative regulator of chondrocyte maturation. Development 133, 651–662 (2006).

54. R. S. Rainbow, H. Kwon, L. Zeng, The role of Nkx3.2 in chondrogenesis. Front Biol (Beijing) 9, 376–381 (2014).

55. Y. Yan, J. Frisén, M. H. Lee, J. Massagué, M. Barbacid, Ablation of the CDK inhibitor p57(Kip2) results in increased apoptosis and delayed differentiation during mouse development. Genes Dev 11 (8):973–83 (1997).

56. N. E. H. Dinesh, N. Baratang, J. Rosseau, R. Mohapatra, L. Li, R. Mahalingam, K. Tiedemann, P. M. Campeau, D. P. Reinhardt, Fibronectin isoforms promote postnatal skeletal development. Matrix Biology 133, 86–102 (2024).

57. G. Luo, P. Ducy, M. D. McKee, G. J. Pinero, E. Loyer, R. R. Behringer, G. Karsenty, Spontaneous calcification of arteries and cartilage in mice lacking matrix GLA protein. Nature 386, 78–81 (1997).

58. A. Farfel, H. Mayan, S. Melnikov, E. J. Holtzman, O. Pinhas-Hamiel, Z. Farfel, Effect of age and affection status on blood pressure, serum potassium and stature in familial hyperkalaemia and hypertension. Nephrology Dialysis Transplantation 26, 1547–1553 (2011).

59. M. Pekkinen, P. A. Terhal, L. D. Botto, P. Henning, R. E. Mäkitie, P. Roschger, A. Jain, M. Kol, M. A. Kjellberg, E. P. Paschalis, K. Van Gassen, M. Murray, P. Bayrak-Toydemir, M. K. Magnusson, J. Jans, M. Kausar, J. C. Carey, P. Somerharju, U. H. Lerner, V. M. Olkkonen, K. Klaushofer, J. C. M. Holthuis, O. Mäkitie, Osteoporosis and skeletal dysplasia caused by pathogenic variants in SGMS2. JCI Insight 4 (7):e126180 (2019).

60. C. W. Stump, The Histogenesis of Bone. J Anat 59 (2):136–154 (1925).

61. A. Cebrián-Silla, C. Alfaro-Cervelló, V. Herranz-Pérez, N. Kaneko, D. H. Park, K. Sawamoto, A. Alvarez-Buylla, D. A. Lim, J. M. García-Verdugo, Unique Organization of the Nuclear Envelope in the Post-natal Quiescent Neural Stem Cells. Stem Cell Reports 9 (1): 203–216 (2017).

62. J. T. Rodgers, K. Y. King, J. O. Brett, M. J. Cromie, G. W. Charville, K. K. Maguire, C. Brunson, N. Mastey, L. Liu, C. R. Tsai, M. A. Goodell, T. A. Rando, MTORC1 controls the adaptive transition of quiescent stem cells from G_0_ to G_Alert_. Nature 510, 393–396 (2014).

63. T. Oichi, J. Kodama, K. Wilson, H. Tian, Y. Imamura Kawasawa, Y. Usami, Y. Oshima, T. Saito, S. Tanaka, M. Iwamoto, S. Otsuru, M. Enomoto-Iwamoto, Nutrient-regulated dynamics of chondroprogenitors in the postnatal murine growth plate. Bone Res 11, 20 (2023).

64. N. Percie du Sert, V. Hurst, A. Ahluwalia, S. Alam, M. T. Avey, M. Baker, W. J. Browne, A. Clark, I. C. Cuthill, U. Dirnagl, M. Emerson, P. Garner, S. T. Holgate, D. W. Howells, N. A. Karp, S. E. Lazic, K. Lidster, C. J. MacCallum, M. Macleod, E. J. Pearl, O. H. Petersen, F. Rawle, P. Reynolds, K. Rooney, E. S. Sena, S. D. Silberberg, T. Steckler, H. Würbel, The ARRIVE guidelines 2.0: Updated guidelines for reporting animal research. PLoS Biol 18 (7):e3000410 (2020).

65. Y. Zhang, G. Parmigiani, W. E. Johnson, ComBat-seq: batch effect adjustment for RNA-seq count data. NAR Genom Bioinform 2, lqaa078 (2020).

66. M. Love, C. Ahlmann-Eltze, K. Forbes, S. Anders, W. Huber, RADIANT EU FP7, NIH NHGRI, CZI, Differential gene expression analysis based on the negative binomial distribution Bioconductor 3.19.

67. G. Marot, A. Rau, F. Jaffrezic, S. Blanck, metaRNASeq: Meta-Analysis of RNA-Seq Data The Comprehensive R Archive Network (2021).

68. M. Kaucka, A. Joven Araus, M. Tesarova, J. D. Currie, J. Boström, M. Kavkova, J. Petersen, Z. Yao, A. Bouchnita, A. Hellander, T. Zikmund, A. Elewa, P. T. Newton, J.-F. Fei, A. S. Chagin, K. Fried, E. M. Tanaka, J. Kaiser, A. Simon, I. Adameyko, Altered developmental programs and oriented cell divisions lead to bulky bones during salamander limb regeneration. Nat Commun 13, 6949 (2022).

69. C.A. Schneider, W.S. Rasband & K.W. Eliceiri, NIH Image to ImageJ: 25 years of image analysis. Nat Methods 9, 671–675 (2012).

70. M. Rabineau, F. Flick, E. Mathieu, A. Tu, B. Senger, J.-C. Voegel, P. Lavalle, P. Schaaf, J.-N. Freund, Y. Haikel, D. Vautier, Cell guidance into quiescent state through chromatin remodeling induced by elastic modulus of substrate. Biomaterials 37, 144–155 (2015).

71. D.C. Barral, L. Staiano, C. Guimas Almeida, D.F. Cutler, E.R. Eden, C.E. Futter, A. Galione, A.R.A Marques, D.L. Medina, G. Napolitano, C. Settembre, V.O Vieira, J.M.F.G. Aerts, P. Atakpa-Adaji, G. Bruno, A. Capuozzo, E. De Leonibus, C. Di Malta, C. Escrevente, A. Esposito, P. Grumati, M.J. Hall, R.O. Teodoro, S.S. Lopes, J.P. Luzio, J. Monfregola, S. Montefusco, F.M. Platt, R. Polishchuck, M. De Risi, I. Sambri, C. Soldati, M.C. Seabra, Current methods to analyze lysosome morphology, positioning, motility and function. Traffic 23 238–269 (2022).

