## Supplementary materials for "Human growth plates house resting zone sub-populations with features of quiescent stem cells"

**Supplementary Materials and Methods**

Collection of mouse tissues

RNAscope *in* situ hybridization and EdU detection

Image capture and analysis specific to supplementary figures

**Supplementary Figures**

Figure S1. Detailed assessment of SRT application to human growth plate biopsies

Figure S2. mRNA in mouse resting zone chondrocytes has predominantly nuclear localization

Figure S3. Low mRNA levels are a strong indicator of LRCs in mice

Figure S4. SRT validation using an additional dataset and *in situ* hybridization

Figure S5. Quantification of sub-populations of human RZ chondrocytes

**Supplementary Table**

Table S1. SRT Validation dataset

### Supplementary Materials and Methods

##### Image capture and analysis

Fluorescent images were acquired using Zeiss laser scanning microscopes (LSM) 700, 800 or 980. Maximal intensity projections (MIP) were routinely generated from the acquired Z-stack images. Depending on the purpose and user, images were analyzed using ImageJ *(69)*, CellProfiler (https://cellprofiler.org/) or QuPath (https://github.com/qupath).

For the quantification of mean poly(A) signal intensity per nucleus as a percentage of maximum value (Figs. 1F, 2I, S2C and S3C), the microscopy MIP files were loaded into ImageJ, and the following steps were used: channels were split, DAPI channel was selected to set its threshold and convert to mask and nuclei was identified. Next, poly(A) channel was selected to measure the mean signal intensity in nuclei. The same settings were used for every sample. The values for each zone were saved as separate csv files and taken into RStudio to analyze. We normalized RNA values within the growth plate of each individual, using its own maximum value (Figs. 1F, 2I, 4D, S2C and S3C). The ggplot package was used to generate graphs (Figs. 1F, 2I, 4D, S2C and S3C). For the quantification of nucleus/cytoplasm poly(A) signal ratio (Figs. 2A, S2D), the microscopy MIP files were loaded into CellProfiler and the following steps were used: multichannel image was split and converted to grayscale, the nucleus was segmented by selecting DAPI channel, the cell was identified by expanding from nucleus edge, the cytoplasm was identified by subtracting nucleus from cell. The poly(A) signal intensity was measured in the nucleus and cytoplasm of each individual cell. The same settings for object size and method were used for every sample. The values for each zone were saved as separate csv files and taken into RStudio to analyze. The ggplot package was used to generate graphs (Figs. 2A, S2D).

For the quantification of triple labeled sections of human growth plate (Fig. 2G), the microscopy MIP files were loaded into ImageJ, and the following steps were used to count the absolute number of labeled cells with BrdU analogues: channels were split, DAPI channel was selected to set its threshold and convert to mask, and nuclei were segmented and counted. Each channel was separately thresholded and labeled nuclei were identified and counted. The data were loaded to GraphPad Prism version 10 to plot and analyze.

For the quantification in Fig. 4, microscopy MIP files were loaded into QuPath and values of mean poly(A) intensity per nucleus obtained following cell detection. A manual assessment of CHRDL2 and SFRP5 signals (in which positive cells either had a large positive region or more than two puncta) was used as a guide to select thresholding values for CHRDL2 and SFRP5 channels. We applied stringent automated thresholding by signal intensity, which could be an underestimate of positive cells, as some cells contained one or two individual puncta, and not all signal was confined to the nucleus.

##### Transmission electron microscopy

A human growth plate slice was further cut during dissection to approximately 1.5 mm in width, fixed in 2.5 % glutaraldehyde (20105; Ladd Research industries) in 0.1 M phosphate (pH 7.4) for 1 hour at room temperature, and stored at 4 °C until further processing (approximately 36 hours). Next, samples were rinsed in 0.1 M phosphate buffer and then post-fixed in 2 % osmium tetroxide (O018; TAAB) /0.1 M phosphate buffer, (pH 7.4) at 4 °C for 2 hours. After stepwise dehydration in ethanol and acetone, the tissue was embedded in LX-112 resin (Ladd). An EM UC7 (Leica) was used to generate Ultrathin sections (approximately 80-100 nm), which were subsequently contrasted with uranyl acetate (U001; TAAB), followed by lead citrate (7398; Merck), and finally examined using a HT7700 transmission electron microscope (Hitachi High-Technologies) at 80 kV. A 2kx2k Veleta CCD camera (Olympus Soft Imaging Solutions) was used to acquire digital images. The proportion of heterochromatin on the nuclear region of electron microscopy images was assessed using ImageJ *(69)* as previously reported *(70)*. In brief, the nuclear membrane was delimited and subjected to an intensity threshold on filtered images by Gaussian blur (radius = 2 pixels). The percentage of heterochromatin is reported as the heterochromatin area in the total nuclear surface. Using electron microscopy images, single-membrane-bound bodies containing electron-dense material with a round- or elongated-like ultrastructure were defined as lysosomes, according to standard criteria *(71)*. The number of lysosomes was assessed by cell.

**Collection of mouse tissues**

Tissues were collected at twenty-eight days of age from wild-type mice on a mixed background. Briefly, hindlimbs were dissected so that the tibia, femur and knee joint were intact, with the skin removed and excess muscle partially trimmed. Samples were fixed in pre-cooled 4 % formaldehyde/PBS for 48 hours at 4°C, then subsequently placed into 30 % sucrose overnight, prior to embedding within OCT medium in cryomolds (62534-1; Tissue-Tek). Samples were stored at -80 °C before processing. In the experiment where LRCs were labeled with EdU (A10044; Invitrogen), mice were given 65 µg EdU/g body mass by daily intraperitoneal injections on post-natal days 6-9.

**RNAscope *in situ* hybridization with EdU detection**

For standard RNAscope *in situ* hybridization, slides were removed from -80 °C freezer and allowed to equilibrate at room temperature until condensation had dried. The samples were incubated in 1x Target Retrieval Solution (322000; ACD) inside a Certoclav CV II/1600 pressure cooker containing 500 ml distilled water, and heated until the pressure reached 0.2 Bar. Slides were then washed in nuclease-free water (NFW [10977-035; Invitrogen]) for 15 seconds, absolute ethanol for 3 minutes, and air-dried. After washing with copious amounts of NFW, sections were covered with Protease III (322337; ACD) and incubated at 40 °C for 30 minutes. The sections were subsequently washed twice in NFW for 2 minutes, incubated with RNAscope Hydrogen Peroxide (322337; ACD) for 10 minutes, and then washed twice by incubating with copious amounts of NFW for 2 minutes. Next, probe hybridization was performed by adding 2-3 drops (~20 µl per drop) of target probes, diluted as per the manufacturer´s instructions, and incubated for 2 hours at 40 °C.

After hybridization, amplification was performed: RNAscope Multiplex v2 Amp1 was applied and incubated for 30 minutes at 40 °C, then RNAscope Multiplex v2 Amp2 was applied and incubated for 30 minutes at 40 °C and finally RNAscope Multiplex v2 Amp3 applied and incubated for 15 minutes at 40 °C. Between all steps, slides were washed twice in 1X Wash Buffer for 2 minutes.

Depending on the number of probes used and their respective channels, the following steps were performed multiple times in sequential order (i.e. from C1-C4) using different fluorophores each time; on the final occasion, DAPI was added to the HRP blocker. RNAscope Multiplex v2 HRP-C1-4 (corresponding to the probes used) were applied and incubated for 15 minutes at 40°C; staining was performed using diluted TSA Plus fluorophores diluted at 1:750 as per the manufacturer´s instructions for 30 minutes at 40°C; HRP blocker (323107; ACD) was applied and incubated for 15 minutes at 40°C. Between all steps, slides were washed twice in 1X Wash Buffer for

2 minutes. After the final wash step, slides were rinsed once in PBS prior to mounting in Fluoroshield (F6182; Sigma-Aldrich) and were stored at 4°C.

When RNAscope was combined with EdU detection, we followed the RNAscope *in situ* hybridization protocol described above, with the following deviations. First, DAPI was not included in the final HRP blocking step, and after the final wash step (mentioned in the paragraph above), we processed the slides as follows: sections were blocked using 3 % normal horse serum in PBS/T (PBS containing 0.05 % Tween20) for 1 hour at room temperature. Sections were washed with PBS/T (3 times for 5 minutes) and exposed to reaction buffer of distilled water containing 100 mM Tris (from 1M stock, pH 7.5), 2 mM $CuSO_4$, 1 µM Alexa-azide 555 (from a 1 mM stock in DMSO), and 100 mM ascorbic acid (added immediately prior to use from a 0.5 M stock in distilled water). Slides were protected from light and incubated at room temperature for 30 minutes, and gently agitated twice during this time. Subsequently, slides were washed once in PBS/T then counterstained with DAPI (1.5 µg/ml in PBS). Slides were then rinsed with PBS (3 times for 5 minutes) and mounted with Fluoroshield.

**Image capture and analysis specific to supplementary figures**

For the quantification of cell number per spot (Fig. S1B), for each zone the number of cells were divided by the area of zone, measured in square micrometre, and multiplied by the area of spots of spatial transcriptomics platform. The values were taken into RStudio to analyze and ggplot package was used to plot.

For the quantification of mean poly(A) signal intensity per nucleus as a percentage of maximum value in EdU-labeled cells in mouse growth plate (Fig. S3C), the microscopy MIP files were loaded into CellProfiler and following steps were used: multichannel image was split and converted to grayscale, each nucleus was identified as an object by selecting DAPI channel, EdU-labeled nucleus was identified as an object by selecting EdU channel, poly(A) signal was measured in nuclei of EdU positive cells and EdU negative cells. The same settings for object size and method were used for every sample. The values for condition were saved as separate csv files and taken into RStudio to analyze. Signal intensity was normalized within the growth plate of each mouse and ggplot package was used to plot.

__Supplementary Figures__

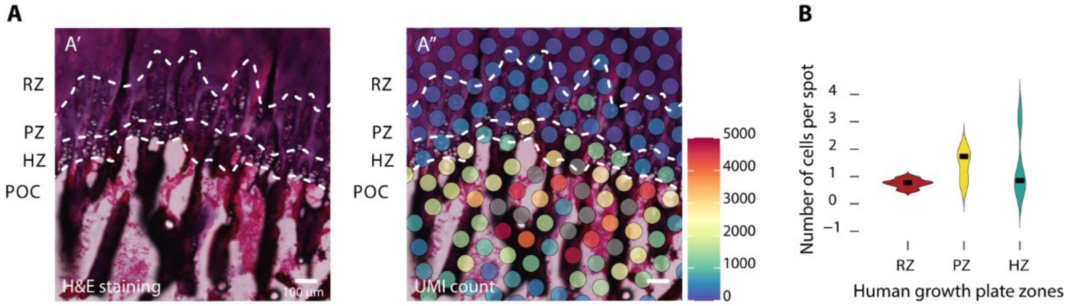


**Figure S1. Detailed assessment of SRT application to human growth plate biopsies.** (**A**) High magnification image indicating bone marrow of the primary spongiosa with hematoxylin and eosin (left) and UMI count overlaid (right). (**B**) Number of cells per 55 $\mu m^2$ (the equivalent area to one Visium spot) was quantified in each zone of human growth plate using DAPI staining of the sections used in panel 1F. The comparison of the number of cells between zones was conducted using Kruskal-Wallis (p-value = 0.1677), the median for each zone is RZ = 0.7834, PZ = 1.7458, HZ = 0.8681 from 5 patients.

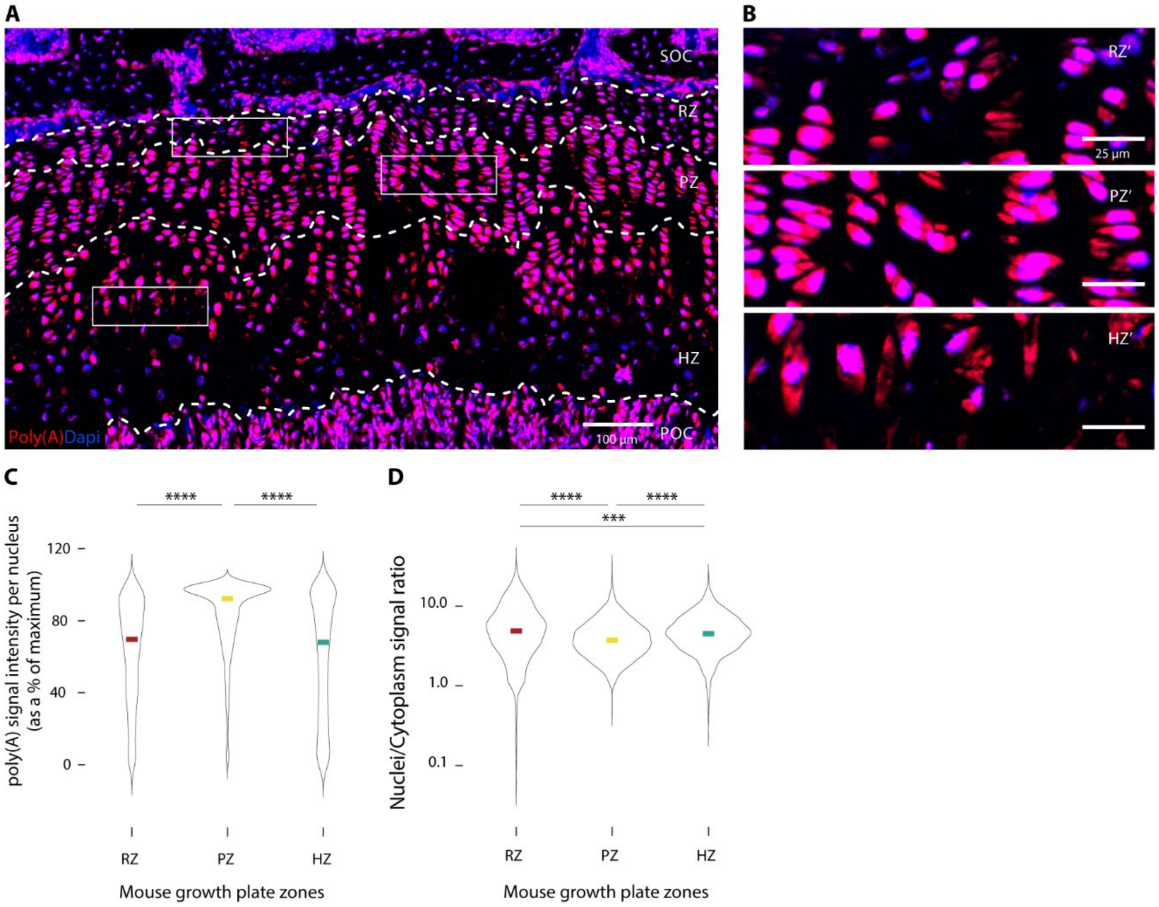


**Figure S2. mRNA in mouse resting zone chondrocytes has predominantly nuclear localization.** (**A**, **B**) RNAscope staining of poly(A) tail of proximal tibial growth plate of mouse at P28. (**C**) Quantification of mean poly(A) signal intensity per nucleus in each growth plate zone. The comparison of medians between zones was conducted using Kruskal-Wallis (p-value < 0.0001) and Wilcoxon post-hoc test: ****p < 0.0001. The median of each zone being RZ = 69.6850, PZ = 92.2472, HZ = 67.9876 (n = 1891 cells in RZ, n = 4197 cells in PZ, n = 2714 cells in HZ). (**D**) RZ chondrocytes of mouse growth plate zones have significantly higher relative poly(A) signal intensity in the nucleus. Data was pooled from 3 mice and analyzed using Kruskal-Wallis (p-value < 0.0001) and Wilcoxon post-hoc test: ****p < 0.0001, ***p < 0.001. The median of each zone was RZ = 4.8801, PZ = 3.7429, HZ = 4.5268 (n = 2657 cells in RZ, n = 6600 cells in PZ, n = 3081 cells in HZ).

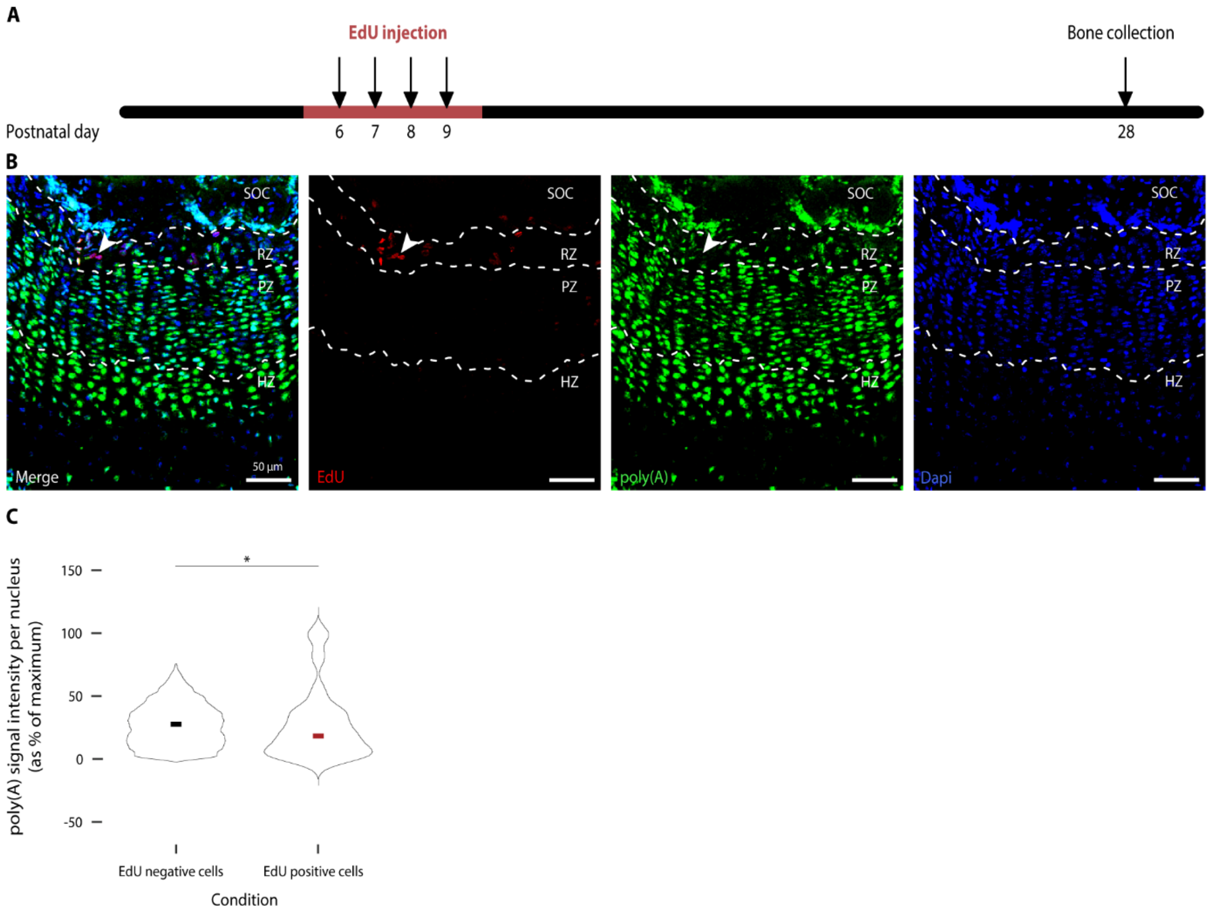


**Figure S3. Low mRNA levels are a strong indicator of LRCs in mice.** (**A**) Workflow for labeling EdU label-retaining RZ chondrocytes in mice at P28. (**B**) Proximal tibial growth plates were stained for EdU and poly(A) tails of RNA using RNAscope. (**C**) Quantification of mean poly(A) signal intensity per nucleus was performed in EdU positive and EdU negative cells. Data was pooled from 4 mice and analyzed using Mann-Whitney-U test: p-value < 0.05. The median of each condition was EdU-negative = 27.70, EdU-positive = 18.49 (n = 6910 for EdU negative cells and n = 45 for EdU positive cells).

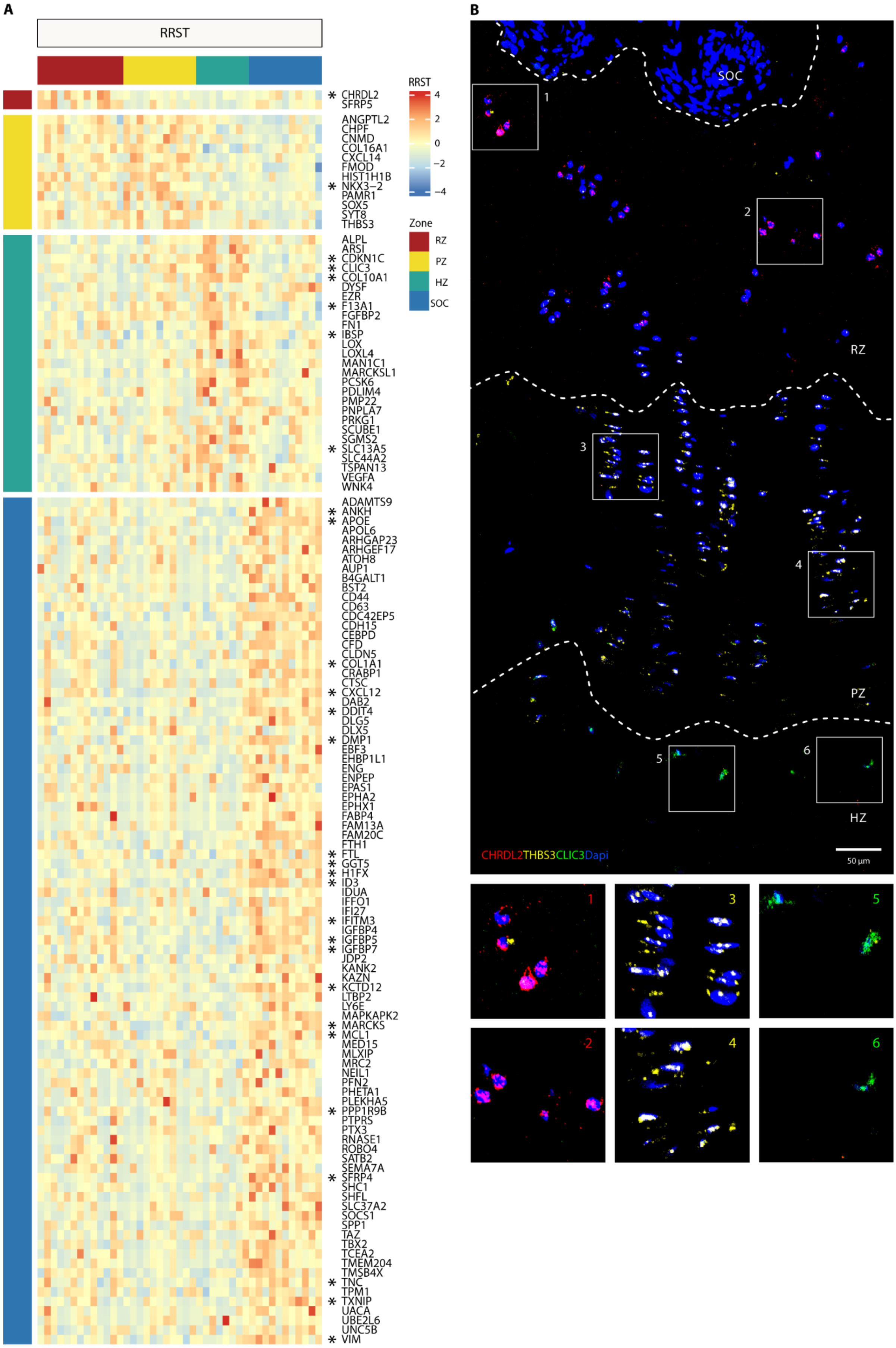


**Figure S4. SRT validation using an additional dataset and *in situ* hybridization.** (A) Expression heatmap of

130 genes significantly upregulated in RRST platform in one of the areas above all others within the validation dataset. Genes labeled with an asterisk were significantly upregulated in the validation dataset. Z-scores for each gene are plotted, blue indicating lower expression and red higher expression of the gene in each sample. The upper bars indicate the sample area, and left bars indicate the area associated with each gene. RZ in red, PZ in yellow, HZ in green and SOC in blue. (**B**) Human growth plate was stained using RNAscope to visualize markers of the RZ (CHRDL2), PZ (THBS3) and HZ (CLIC3) identified by SRT; n = 1 patient.

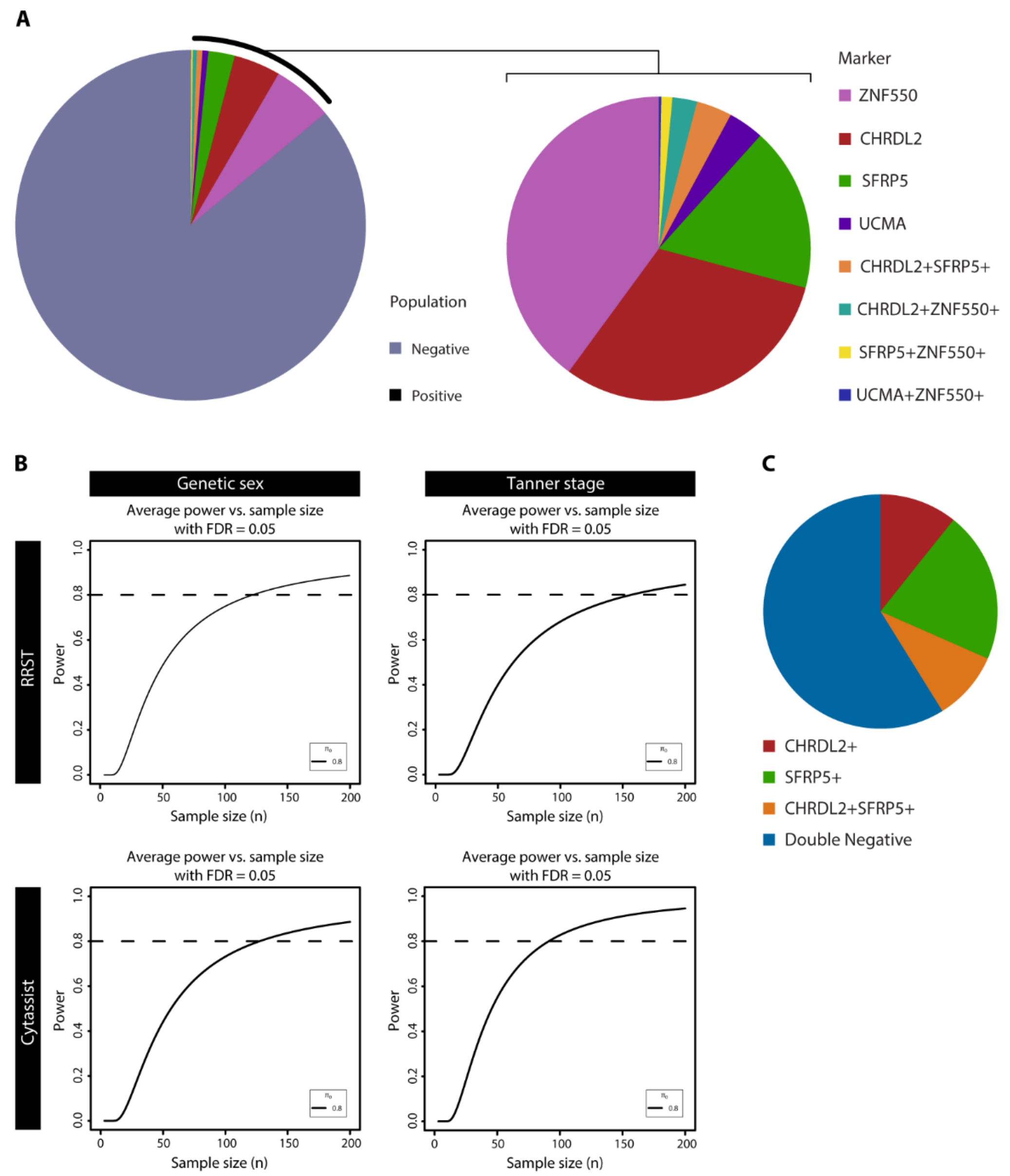


**Figure S5. Quantification of sub-populations of human RZ chondrocytes.** (**A**) Pie charts visualizing the relative abundance of spots containing the various sub-populations of RZ markers identified from SRT from the entire population (left) and among sub-populations (right). (**B**) Power analysis was performed to determine the number of patients needed to be included to have a power of 0.8 for genetic sex and Tanner stage, based on both platforms. (**C**) Pie chart visualizing the relative abundance of spots containing sub-populations of RZ markers identified by their expression of CHRDL2 or SFRP5 using RNAscope.

**Supplementary Table**

| Patient ID | Platform | Genetic sex | Tanner pubertal stage PH | Tanner pubertal stage B | Tanner pubertal stage G | Patient age at surgery |
|---|---|---|---|---|---|---|
| 1 | RRST | M | 3 | - | 3 | 13 y 2 m |
| 1 | RRST | M | 3 | - | 3 | 13 y 2 m |
| 4 | RRST | F | 3 to 4 | 4 | - | 12 y 11 m |
| 5 | RRST | M | 2 | - | 2 | 12 y 2 m |
| 6 | RRST | F | 4 | 4 | - | 14 y 6 m |
| 6 | RRST | F | 4 | 4 | - | 14 y 6 m |
| 9 | RRST | M | 4 | - | 4 | 13 y 10 m |
| 10 | RRST | M | 4 | - | 5 | 14 y 4 m |

**Table S1. SRT Validation dataset.** Metadata of the patient information regarding the tissues analyzed in Fig. S4A. Patient IDs numbering correspond to a continuation from Table 1. Duplication in this table indicates samples from the same patient were run in different capture areas.
